# Elements of a stochastic 3D prediction engine in larval zebrafish prey capture

**DOI:** 10.1101/755777

**Authors:** Andrew D Bolton, Martin Haesemeyer, Josua Jordi, Ulrich Schaechtle, Feras Saad, Vikash K Mansinghka, Joshua B Tenenbaum, Florian Engert

## Abstract

Many predatory animals rely on accurate sensory perception, predictive models, and precise pursuits to catch moving prey. Larval zebrafish intercept paramecia during their hunting behavior, but the precise trajectories of their prey have never been recorded in relation to fish movements in three dimensions.

As a means of uncovering what a simple organism understands about its physical world, we have constructed a 3D-imaging setup to simultaneously record the behavior of larval zebrafish, as well as their moving prey, during hunting. We show that zebrafish robustly transform their 3D displacement and rotation according to the position of their prey while modulating both of these variables depending on prey velocity. This is true for both azimuth and altitude, but particulars of the hunting algorithm in the two planes are slightly different to accommodate an asymmetric strike zone. We show that the combination of position and velocity perception provides the fish with a preferred future positional estimate, indicating an ability to project trajectories forward in time. Using computational models, we show that this projection ability is critical for prey capture efficiency and success. Further, we demonstrate that fish use a graded stochasticity algorithm where the variance around the mean result of each swim scales with distance from the target. Notably, this strategy provides the animal with a considerable improvement over equivalent noise-free strategies.

In sum, our quantitative and probabilistic modeling shows that zebrafish are equipped with a stochastic recursive algorithm that embodies an implicit predictive model of the world. This algorithm, built by a simple set of behavioral rules, allows the fish to optimize their hunting strategy in a naturalistic three-dimensional environment.

## INTRODUCTION

Predatory animals ranging from dragonflies to frogs, salamanders and falcons possess internal models of prey motion. These models allow predators to extrapolate the path of a moving prey item and predict future locations of their prey (Mischiati et al. 2015, Arbib 1987, Colett 1982, Borghuis and Leonardo 2015, Brighton et al. 2017). Prey capture behaviors therefore reflect “physical knowledge” that allows the animal to thrive in an environment where concepts of 3D space, object position, and motion must be leveraged for survival.

While much of the physical knowledge displayed by animals and humans is indeed complex (Lake et al. 2017, Battaglia et al. 2013, Ullman et al 2017, Spelke and Hespos 2001, Baillargeon 1987, Johnson et al. 2003), it is becoming clear from recent ethological and neuroscience studies that remarkable capabilities in animals are often built by combining sets of more basic behaviors. For example, the theory of Self-Organization states that seemingly complicated behaviors like schooling in fish arise from relatively simple visuomotor rules being executed by members of the group, without reference to the emerging global pattern (Couzin and Krause 2003). Bees learning to pull strings for hidden rewards appear to be displaying insight and ingenuity, but are in fact instituting a set of observational and associative learning rules in sequence (Alem et al. 2016). The synthesis of complexity from more basic behavioral modules has long been a staple of computer science and artificial intelligence (Brooks 1991); Marvin Minsky proposed that intelligence is brought about by the interaction of mindless “agents” in his Society of Mind theory (Minsky 1986), while life-like goal oriented behavior was similarly formulated in Braitenberg’s “Vehicles” (Braitenberg 1984).

The construction of sophisticated behaviors from simple rules is also observed in the larval zebrafish, a teleost with 100,000 neurons that has established itself as a tractable vertebrate model in modern neuroscience. In zebrafish, a seemingly complex behavior like the stabilization of position in turbulent streams can be accomplished by instantiating a curl-detector for local water flow and an optomotor response, where whole field visual motion leads to stereotypic swimming and turning behaviors (Oteiza et al. 2017, Naumann et al. 2016). Moreover, a strategy of energy-efficient foraging emerges from the zebrafish’s innate tendency to locomote via alternating series of unidirectional turns (Dunn et al. 2016).

Interestingly, most zebrafish behaviors, including their locomotor and optomotor schemes, are actually probabilistic in nature. For example, the precise number of unidirectional turns in any spontaneous swimming stretch is stochastic, while turn magnitude in response to angular optic flow varies widely (Dunn et al. 2016, Naumann et al. 2016). Noise in *any* computing system is usually considered inconvenient and a nuisance to be overcome (Kording and Wolpert 2012). However, there is precedent for noisy sensory detection and stochastic movements working to the benefit of many animals. Crayfish and paddlefish, for instance, both take advantage of stochastic resonance to detect sparse prey and predators (Douglass et al. 1993, Russell et al. 1999). Beneficially stochastic foraging has been observed in animals ranging down to micro-organisms, while predated animals used mixed stochastic strategies to avoid predictability by predators (Jensen 2018). Neurons themselves benefit from gaussian noise injection, which actually improves detection of a periodic stimulus (Wilson 1999). Nevertheless, it is unknown whether the stochasticity observed in zebrafish behavior is also beneficial to the animal.

In this study, we addressed whether the generation of complex, efficient behavior from simple stochastic rules also applies to the most complicated behavior that larval zebrafish perform: prey capture. Prey capture requires the fish to select, pursue, and ultimately consume fast moving single-celled organisms swimming through their environment. Elegant work has been performed in documenting the types of body movements the fish use during prey capture and in the characterization of neurons responsible for prey perception (Johnson et al. 2019, Marques et al. 2018, Gahtan et al. 2005, Bianco and Engert 2015, Muto et al. 2013). Furthermore, foundational studies have led to an initial understanding of how fish choose their movements based on prey contingencies (Patterson et al. 2013, Trivedi and Bollmann 2013, Bianco et al. 2011). We build upon these studies by analyzing this behavior in its natural 3D setting, whereas previous studies have typically neglected vertical fish and prey movements. Moreover, we develop an experimental and computational framework that can simultaneously record fish and prey trajectories. This approach allowed us to accurately map the fish’s sensorimotor transformations in response to ongoing prey features, which are described in an egocentric spherical coordinate system that reflects the fish’s three-dimensional point of view. Particularly, we illustrate three main rules governing prey pursuit. First, sensorimotor transformations during prey capture are largely controlled by the azimuth angle, altitude angle and computed radial distance of prey before the fish initiates a pursuit movement. Second, all aspects of the fish’s 3D movement choices are strongly and proportionally modulated by the angular and radial velocity of its prey. Combining these two rules yields an emergent strategy whereby the fish can predict future prey locations and recursively maneuver prey to a preferred post-movement 3D position. Third, we show that the speed of the fish’s recursive hunting strategy is benefited by noise in its sensorimotor transformations. Importantly, this stochasticity is graded, meaning that the further away a prey item is from the fish’s goal, the more variable the outcome of the fish’s choice becomes. Using a series of computational models and virtual prey capture simulations, we show that position perception, velocity projection, and graded variance are in fact *all* essential for effective and energy-efficient prey capture performance.

This work reveals that even the most complex behavior in larval zebrafish can be reduced to a set of simple rules. These rules coalesce to generate a stochastic recursive algorithm embodied by zebrafish during hunting, which ultimately reflects an implicit predictive model of the world.

## RESULTS

### Developing A 3D Environment For Elucidating Hunting Sequences In Zebrafish

We first sought to characterize the sensorimotor transformations larval zebrafish implement when pursuing and capturing paramecia. Hunting sequences were recorded from 46 larval zebrafish using a behavioral setup that could simultaneously image the fish and its prey from the top and side at high resolution and speed (Figure 1A). Custom computer vision software was designed to reconstruct the fish’s 3D position and two of the principal axes, yaw and pitch (Figure 1B), as well as the trajectories of paramecia in the environment (Figure 1C). These reconstructions allowed us to spatially map prey position and velocity to an egocentric spherical coordinate system originating at the mouth of the fish. Hereby each paramecium is assigned an azimuth, altitude and distance as a positional coordinate (“Prey Az, Prey Alt, Prey Dist”), along with angular (Prey δAz / δt, Prey δAlt / δt) and radial (Prey δDist / δt) velocities with respect to the fish’s 3D point of view (Figure 1D). To view the prey environment from the fish’s reconstructed 3D perspective, see Supplemental Video 1.

**Figure 1:**
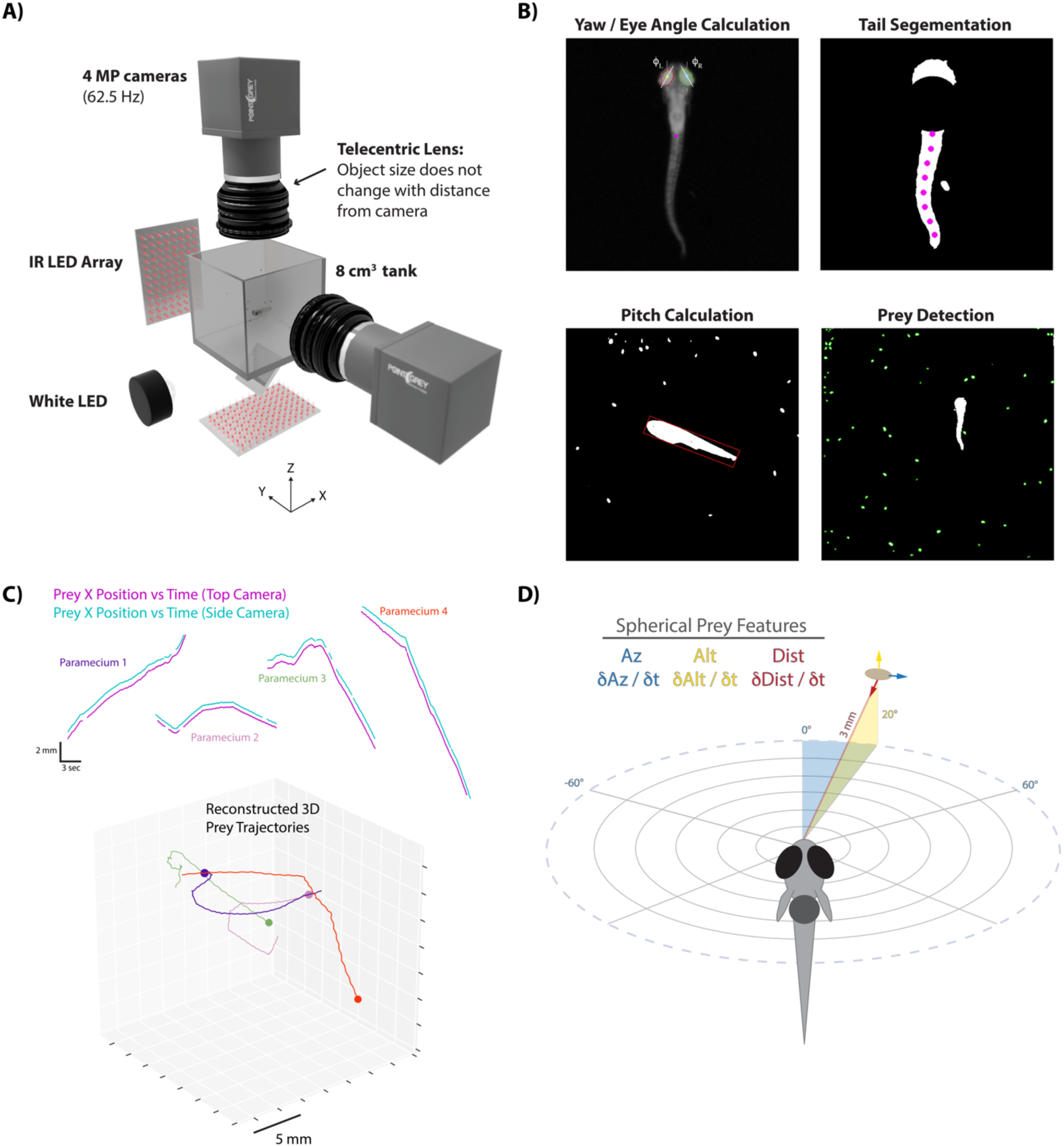
A) 3D rendering of rig design and features. B) Computer vision algorithms extract the continuous eye angle, yaw, pitch, and tail angle of the zebrafish. In every frame prey are detected using a contour finding algorithm. C) Prey contours from the two cameras are matched in time using a correlation and 3D distance-based algorithm, allowing 3D reconstruction of prey trajectories. D) Prey features are mapped to a spherical coordinate system originating at the fish’s mouth. Altitude is positive above the fish, negative below. Azimuth is positive right of the fish, negative left. Distance is the magnitude of the vector pointing from the fish’s mouth to the prey.

Larval zebrafish swim in discrete “bouts”, which consist of a pulse of velocity lasting ∼200 ms, followed by a variable period of intermittent quiescence (Figure 2A; Budick and O’Malley 2000). We took advantage of this unique facet of fish behavior to frame each bout performed during a hunting sequence as an individual “decision” based on the spherical position and velocity of pursued prey. Specifically, we identified the start-time and end-time of each swim bout using fluctuations in tail variance and velocity (see Methods). This allowed us to precisely understand how the fish transforms pre-bout prey features into movements that displace and rotate the fish to a new location in 3D space at the end of the bout.

**Figure 2:**
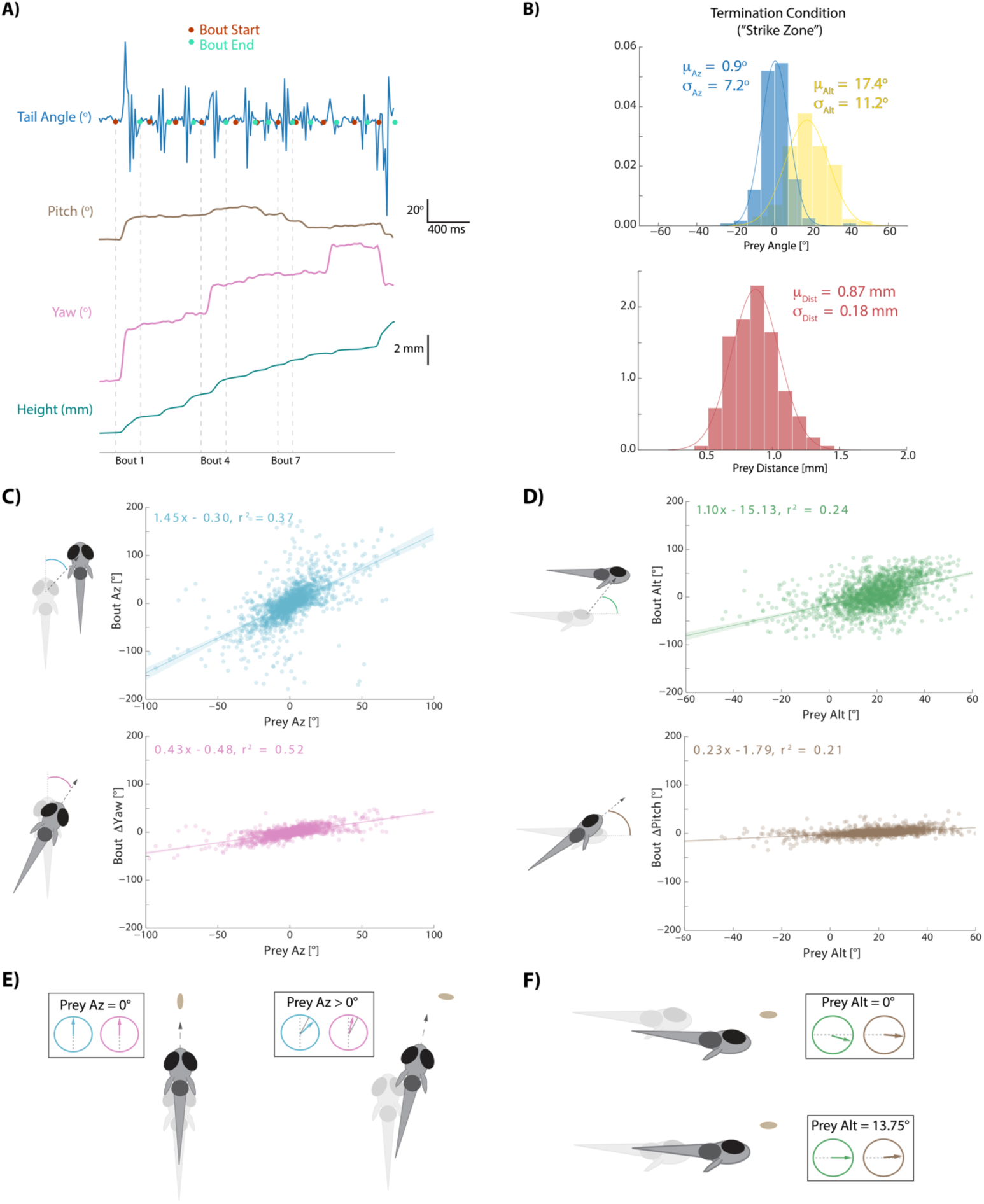
A) During hunt sequences, fish swim in bouts that can be detected using tail variance. Bouts can change the yaw, pitch, and position of the fish, while time between bouts is marked by quiescence. B) Histograms showing the distributions of spherical prey positions when fish successfully ate a paramecium during a strike. C, D) Regression fits between prey position and bout variables executed by the fish. E, F) Features of sensorimotor transformations based on prey position: fish swim forward if prey are directly in front. Otherwise, if prey are on right, fish displace and rotate right; and vice versa. Fish displace downward if prey are at 0° altitude, but displace with no altitude change if prey are at 13.75°. In all schematics (C-F), positions and orientations at the beginning of the bout are represented by transparent fish, and by opaque fish at the end of the bout.

Hunting sequences themselves consist of an initiation bout, multiple pursuit bouts, and a termination bout. Initiation bouts were identified by detecting whether the eyes on a given bout have converged. Eye convergence, which allows the use of stereovision by creating a small binocular zone, is a well-known correlate of hunting state entrance in zebrafish (Gahtan et al. 2005, Bianco et al. 2011). Hunt sequences were therefore identified by clustering the continuous eye angle record for both eyes during each bout (Supp Fig 1A). Hunt sequences were terminated on the bouts where fish either struck at their pursued prey or clearly quit pursuit. Quitting has been called an “aborted” hunt in the literature (Henriques et al. 2019, Johnson et al. 2019), and most aborts in our dataset, as in other studies, corresponded to the cluster demarking deconvergence of the eyes (Supp Fig 1B) and a return to an exploratory state. The average hunting sequence ending in a strike in our dataset lasted for 5 bouts (IQR = 4 to 7).

### Zebrafish Typically Choose The Closest Prey Item When Initiating A Hunt Sequence

Nearly all hunt sequences in our dataset began with the choice of a single prey item to pursue (Supp Fig 1B). The choice of prey item was straightforward. Fish almost invariably chose the closest paramecium in the environment conditioned on the fact that the paramecium was fairly close to its midline in azimuth and significantly above it in altitude (Supp Fig 2A, 2C; μ_az_ = 0.7° σ_az_ = 48.4°; μ_alt_ = 19.9° σ_alt_ = 19.7°; μ_dist_ = 3.4 mm σ_dist_ = 1.6 mm). There was no particular bias of prey choice in terms of direction or magnitude of velocity (Supp Fig 2B).

### Sensorimotor Transformations During Prey Capture Are Largely Controlled By Pre-Bout Prey Position

After choosing a prey item during an initiation bout, the fish engages in a series of pursuit bouts (see Supplementary Video 1) that can each influence the position, yaw, and pitch of the animal (Figure 2A). Pursuit bouts are conducted until prey are positioned in a “strike zone”, which defines the termination condition for successful hunts in spherical coordinates relative to the fish: this zone is directly in front of and considerably above the fish (Figure 2B; avg. 0.9° Prey Az, 17.4° Prey Alt, .870 mm Prey Dist; Mearns et al. 2019). We investigated whether displacements and rotations during pursuit bouts were influenced by 3D prey position before each bout. For the rest of this manuscript, sensorimotor transformations during hunt sequences in which a strike was performed are described. However, the algorithm the fish uses during aborts is nearly identical (Supp Fig 3; note that the last 3 bouts in abort sequences tend to go awry for unknown reasons related to nucleus isthmi activity; Henriques et al. 2019).

For every pursuit bout, we calculate an axis of motion in egocentric spherical coordinates along which fish displace during the bout. This axis is defined by an azimuth and an altitude angle (“Bout Az”, “Bout Alt”), and the magnitude of displacement along this axis is termed “Bout Distance”. Because the axis of motion during bouts is not perfectly aligned to the axis of symmetry, yaw and pitch changes are independently described per bout (“Bout ΔYaw”, “Bout ΔPitch”). Diagrams of rotation and displacement variables are provided alongside Figures 2C and 2D.

Each bout aspect is primarily controlled by the position of the prey relative to the fish immediately anteceding bout initiation (Figure 2C, 2D). Regression fits show that Bout Az and Bout ΔYaw are well correlated to Prey Az, with negligible offset (0.3° and 0.48°). A similar linear relationship is seen when mapping Bout Alt and Bout ΔPitch of pursuit bouts to Prey Alt, but this time with significant negative offsets (−15.13° and −1.79°). These negative offsets imply that if Prey Alt before a pursuit bout is 0°, which one may preconceive as the fish’s ultimate “goal”, the fish will dive downwards by ∼15° and rotate downward by ∼2°. Overall, the regression fits in Figure 2C and D imply that the fish generally swims straight forward if prey are directly in front of them (∼0° azimuth, Figure 2E), and otherwise the positive Bout Az and Bout ΔYaw slopes establish that if the prey is on the right, the fish rotates and displaces proportionally to the right. If the prey is on the left, vice versa. This proportional relationship is similar in altitude, but due to the negative offsets in Bout Alt and Bout ΔPitch, inflection points are not at ∼0°, but at 13.75° and 7.8° respectively. This negative bias serves to consistently maintain the prey above the fish at the end of pursuit bouts. Schematics of these rules are shown in Figure 2E and 2F. Interestingly, these positional transformations reflect the fish’s preferred position in which to strike for prey consumption: directly in front and significantly above (Figure 2B).

Bout Distance along the axis of motion established by Bout Az and Bout Alt is a more complex variable and will be addressed below.

### All 3D Movements During Prey Capture Are Strongly Modulated By Prey Velocity

Previous studies had suggested that kinematics of zebrafish movements change from slow to fast bouts based on whether prey are approaching or swimming away from the fish (Patterson et al. 2013). Our 3D setup allowed us to conduct a detailed analysis of prey velocity perception in all planes. We find that every movement the fish performs during prey capture is strongly and proportionally influenced by both the radial and angular velocity of its prey. This is likely because the prey move very quickly around the fish in our assay (74% 3D vector velocity > 3 paramecium lengths per second, 27% > 6 paramecium lengths per second; avg. Prey δAz / δt = 29° / s, Prey δAlt / δt = 25° / s for pursuit bouts leading to strikes; Figure 3A). An example of bout feature modulation by prey velocity is depicted in Figure 3B and 3C. When prey are in the window between 0° and 5° in azimuth (i.e. slightly right of the fish), fish will rotate and displace to the right if the prey is moving away from the fish, but on average to the left if the prey is moving towards them, anticipating that the prey will cross its midline. Likewise, in the altitude window 15°-20° above the fish, fish most often “wait” for the prey to arrive if prey are moving downward (Bout Alt ∼0°) while showing characteristic upward displacement and pitch if prey are moving upwards in altitude.

**Figure 3:**
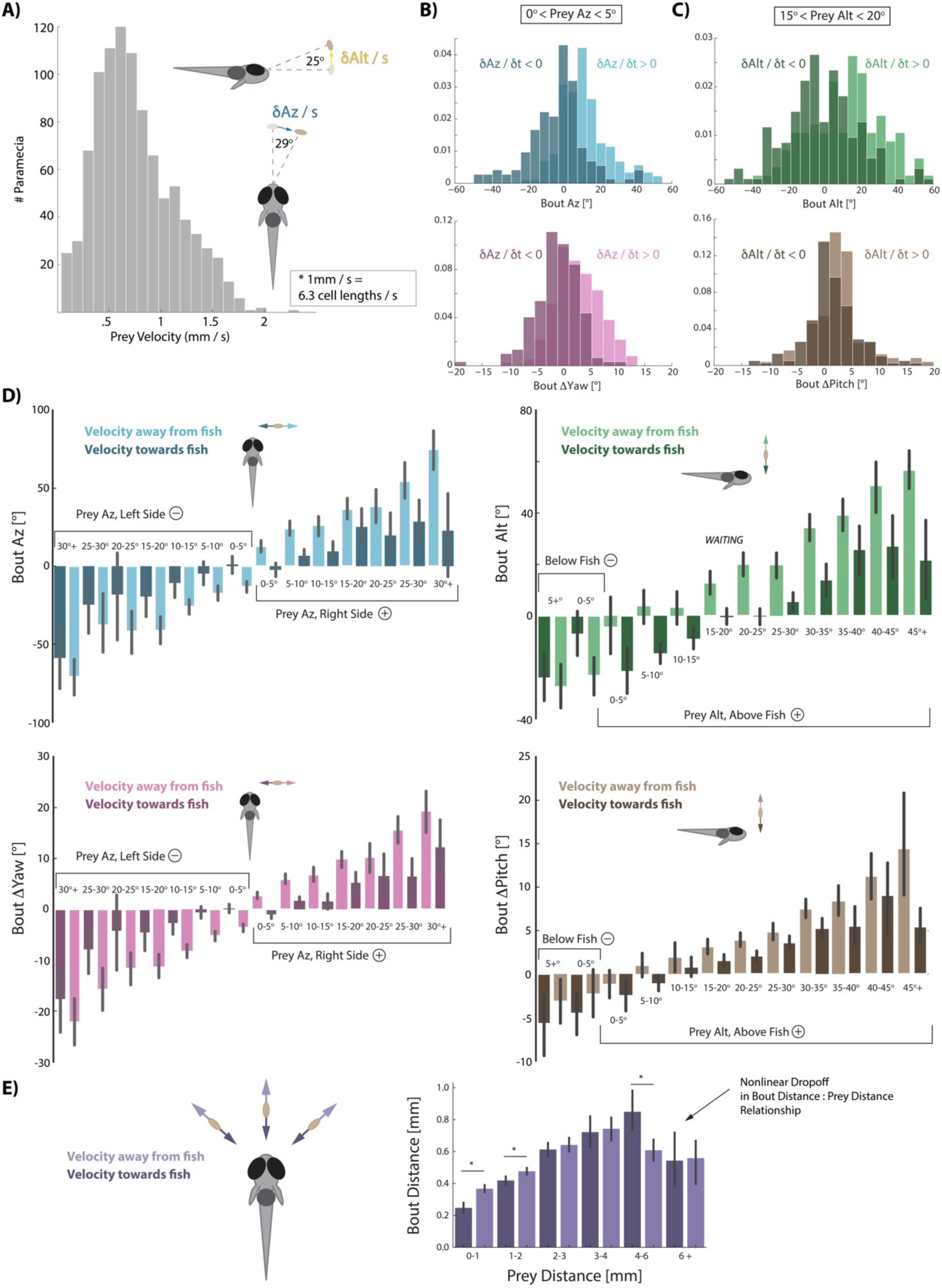
A) Distribution of 3D velocities of prey hunted by fish in our study. Mean azimuth and altitude velocities (absolute value) averaged over 80 ms before pursuit bout initiation, which is the time interval of all velocity calculations implemented here. B, C) Light colors show distributions of each bout variable if prey are moving away from the fish while dark colors show distributions if prey are moving towards the fish in the same 5° window of space. Velocity significantly separates the distributions, which occurs in all 5° spatial windows across all variables (D). A pattern emerges of dampening movements when prey are moving towards and amplifying movements when prey are moving away. Multiple regression fitting of bout variables to prey position and velocity in all planes confirm and quantify the dampening and amplification (azimuth velocity moving left to right of the fish is positive, altitude velocity upward is positive). Bout Az is biased by .*251* * *Prey* δAz / δt (.219 - .283 95% CI), Bout ΔYaw by .*054* * *Prey* δAz / δt (.046-.061 95% CI), Bout Alt by .*300* * *Prey* δAlt / δt (.268 - .331 95% CI), and Bout ΔPitch by .*031* * *Prey* δAlt / δt (.024 - .039 95% CI). E) Bout Distance is linearly proportional to Prey Distance but only within 4 mm of the fish. Likewise, dampening of Bout Distance when Prey δDist / δt < 0 (prey approaching radially), and amplifying when Prey δDist / δt > 0 (prey moving afar), occurs in two windows: 0-1 mm, and 1-2 mm from the fish. Overall, however, multiple regression finds: *Bout Distance =* .*105* * *Prey Dist +* .*053* * *Prey* δDist / δt (95% CIs = .094-.116, .034-.071), which reflects the fact that ∼70% of pursuit bouts occur when prey are within 2 mm.

Indeed, averages of all bout variables in every 5° window of prey space are modulated by the direction of prey velocity (Figure 3D); all bout variables are amplified if prey are moving away from the fish and dampened when prey are moving towards the fish. This modulation of bout features is proportional to the velocity of the prey, as it is well fit by multiple linear regression transforming position *and* velocity of prey into fish bout variables (see Figure 3D legend). Provided that most bouts zebrafish make during hunts are 176 ms long (IQR = 144 ms to 208 ms), we projected the velocity regression coefficients in Figure 3D forward in time by dividing the average bout length. This yields coefficients transforming **projected** prey position change by 1.43 for Bout Az, .31 for Bout ΔYaw, 1.70 for Bout Alt, and .18 for Bout ΔPitch. Critically, these values closely approximate the coefficients describing bout transformations to prey position itself in Figure 2; this is the first hint that the fish has reduced the problem of position prediction to adding velocity multiplied by bout time to its current prey position percept. In this sense, the fish are performing Euler’s Method of approximating a future position based on its instantaneous derivative.

The final bout variable to address, Bout Distance, is more nuanced than the other four bout features. The linear relationship between Bout Distance to Prey Distance, in general, is only strong when prey are < 4mm from the fish (Figure 3E). Bout Distance is significantly modulated by radial prey velocity (δDist / δt; see Figure 1D) when prey are within 2 mm (Figure 3E). In this spatial window, radial velocity of prey coming towards the fish dampens Bout Distance while radial velocity moving afar from the fish amplifies Bout Distance. Multiple regression finds an overall correlation between Bout Distance and Prey Distance with significant dampening when Prey δDist / δt < 0 and amplification of Bout Distance when Prey δDist / δt > 0 (Figure 3E legend for coefficients).

### Computational Models Of Prey Capture Behavior Show Efficiency And Success Arise From Velocity Perception

Next, we asked to which degree the use of prey velocity information contributes to the animal’s ability to efficiently and successfully capture prey. To that end we constructed a Virtual Prey Capture Simulation Environment in which computational models of fish behavior with different “powers” can be pitted against each other. We started by re-creating the initial conditions of 225 hunt sequences initiated by the fish that resulted in a real-life strike (Figure 4A, Methods). From these initial conditions we launched five models that transform current paramecium features into 3D bouts as the prey moves through the environment. Each model initiates a bout at the precise moment when the real fish initiated a bout during the sequence, and model sequences are terminated when the prey enters the virtual strike zone (Figure 2B, Methods). Moreover, we compare all models to the real 3D fish trajectory (the “real fish” model, Figure 4 Model 1 [Blue]), both assuring that our 3D bout and strike-zone characterizations recapitulate real-life performance, and allowing us to compare post-bout coordinates of all models to the characterization of real bouts.

**Figure 4:**
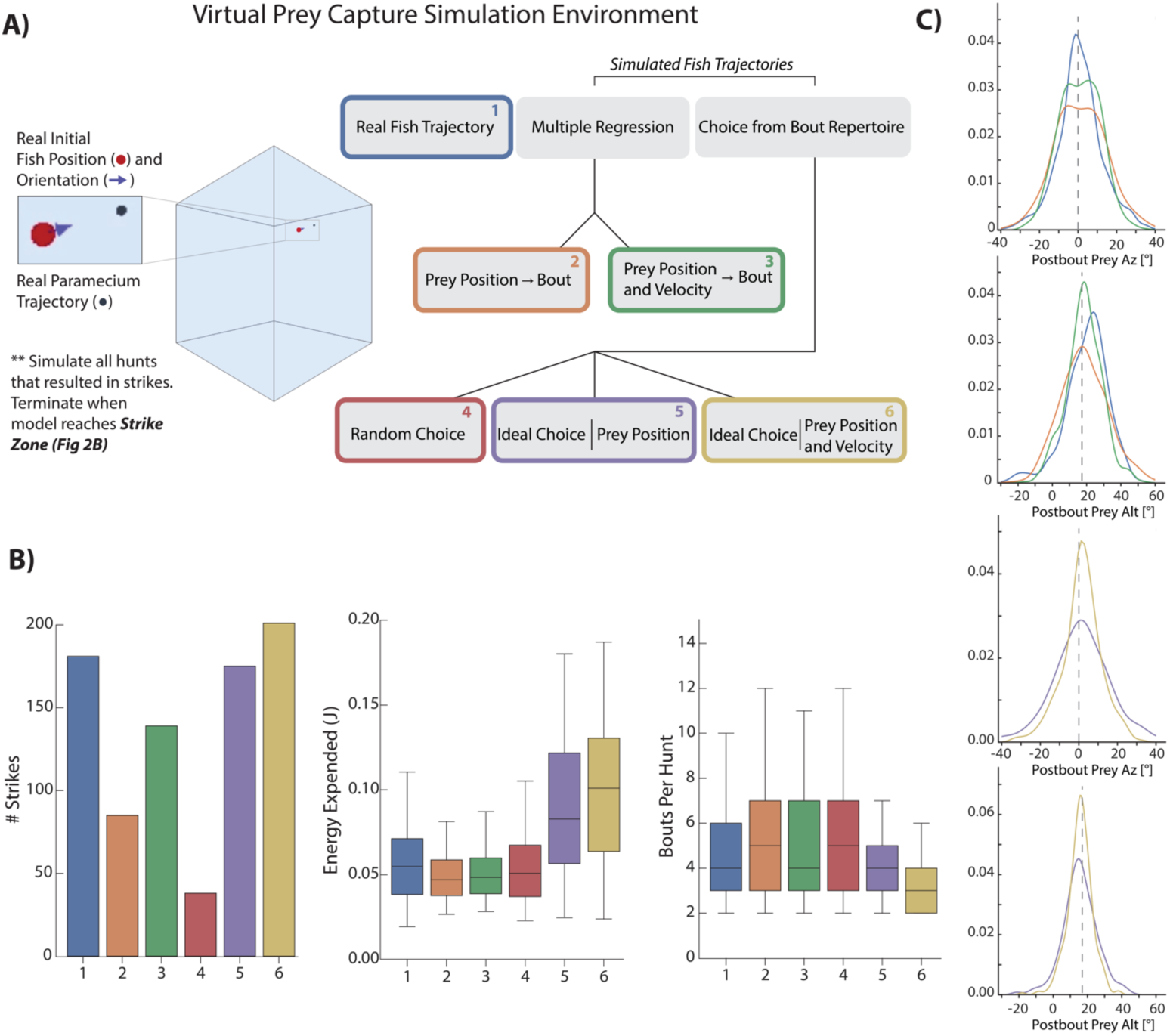
A) A virtual prey capture environment mimicking the prey capture tank was generated to test 3 different types of models, and 6 models overall, described in the schematic. Models control a virtual fish consisting of a 3D position (red dot) and a 3D unit vector pointing in the direction of the fish’s heading. Virtual fish are started at the exact position and rotation where fish initiated hunts in the dataset. Prey trajectories are launched that reconstruct the real paramecium movements that occurred during hunts. The virtual fish moves in bouts timed to real life bouts, and if the prey enters the strike zone (defined by the distributions in Figure 2B, Methods), the hunt is terminated. B) Barplots showing performance of all 6 models in success (# Strikes), energy use per hunt sequence, and how many bouts each model performed during the hunt (a metric of hunt speed). C) KDE plots showing the distribution of Post-Bout Prey Az and Post-Bout Prey Alt distributions for each model during virtual hunts. Dotted lines demark the strike zone mean.

The capabilities of each model are as follows (Figure 4A): First, a multiple regression model was fit on only positional features of the paramecium (Model 2 [Orange]). This model linearly transforms current position of the prey into 3D bout features according to the regression fits in Figure 2C, D (and the bout distance fit from 3E). Model 3 (Green), a multiple regression model fit on both the position and velocity of prey, accounts for the amplification and dampening of bout features by prey velocity in Figure 3; position transforms are thus linearly biased by the velocity coefficients described in the Figure 3D and 3E legend.

Models 4-6 are “Choice” models which can draw from a distribution of 1782 pursuit bouts conducted by fish in our dataset during sequences ending in a strike. This bout pool can be thought of as the “pursuit repertoire” of larval zebrafish. Model 4 (Red) simply draws a random bout from the bout pool at every juncture. Model 5 (Purple) chooses the ideal bout from the pool, with each bout scored on its achievement in reducing the prey’s azimuth, altitude, and distance to the mean values of the strike zone (see Methods). Model 5 does *not* have access to the velocity of the prey, meaning that it will zero in on the *pre-bout* prey position. Lastly, Model 6 (Gold) has the same capabilities to choose ideally as Model 5, but will extrapolate the current prey velocity and add its time multiplied bias to the current position. Therefore, Model 6 can predict the future paramecium position at the end of the bout, but choose ideally instead of linearly.

To compare the models, we describe four facets of their performance intended to score raw hunting success as well as energetic cost: First, how many times out of 225 the model achieved success in placing the virtual prey in its strike zone. Second, how much total energy was expended by the bout combination used during the hunt (see Methods). Third, how many total bout choices the model made per hunt as a metric for capture speed (Figure 4B). And lastly, the Post-Bout Prey Az and Alt prey coordinate for each transformation was plotted to illustrate how well each model does in reducing prey coordinates to the strike zone (Figure 4C). Performance of each model for a given prey trajectory can be viewed in Supplemental Video 2.

The first clear result is that the velocity-based regression Model 3 (Green) improves hunting success over the position-only regression Model 2 (Orange) by 65%. Moreover, the average number of bouts is one less for the velocity model, matching the average of the real fish (Blue). This indicates that the velocity information processed by the fish, which allows projection of future prey coordinates, is critical for both its success rate and speed in capture. When examining the Post-Bout Prey Az for the regression models, the velocity-based Model 3 shows a tighter distribution around 0°, indicating that it is closer to the strike zone on average than position-only Model 2 (Figure 4C, right top panels). This is also true for Post-Bout Prey Alt, where the green plot shows a stronger bias towards the strike zone.

Velocity information is also critical to the performance of ideal choice models. As expected, the random choice Model 4 (Red) performs very poorly, indicating that although similar “types” of pursuit bouts are chosen, success can only be gained by accounting for prey features. Model 5 (Purple), interestingly, does not outperform the real fish in terms of success (3% worse) or average speed of capture, and expends significantly more energy. Model 5 therefore issues high energy bouts (i.e. bouts that strongly rotate and displace the fish), but without any average improvement over what the fish actually did, owing to the high velocity of the average prey item (Figure 3A). Model 6 (Gold), however, by accounting for prey velocity and choosing ideally, improves success rate over Model 1 (Real Fish) and 5 (Ideal Position) by 14% and 17%, reduces the average number of required bouts by 1, and more effectively reduces Post-Bout Prey Az and Alt to the strike-zone (Figure 4C, bottom panels). Nevertheless, Model 6 expends the most energy of any model per hunt sequence, meaning that fully ideal choice comes at an energetic cost. We therefore conclude that position perception improves performance over issuing random pursuit bouts with no reference to the prey, and that velocity information in all formats improves model performance over position perception alone. Of note, the high energy usage of the ideal models relative to the real fish argues against the natural implementations of these strategies. Lastly, although the real fish takes fewer bouts to reach the target than the regression models (#2, #3), it requires slightly more total energy to do so. This implies that a modicum of additional energy is expended per bout, and we speculate that the generation of stochasticity in the real fish’s algorithm (described below) is to blame.

### Pre-Bout To Post-Bout Prey Coordinate Transformation Reveals A Canonical Hunting Strategy

We next examined how the fish’s sensorimotor transformations described in Figure 2 and 3 convert pre-bout coordinates of prey into more favorable coordinates after bout completion. This step to a higher level of abstraction in characterizing prey capture allowed us to consider the fish’s ultimate objectives, getting us closer to a general description of capture strategy.

In Figure 5A, light colors show these transformations when the prey is moving away from the fish while dark colors represent when the prey is moving towards the fish. Regardless of velocity direction, each bout reduces Prey Az as well as Prey Alt on average by ∼.5, maintaining the small bias (∼8°) in Prey Alt observed in Figure 2. These data imply that the fish’s strategy is to reduce prey angle in both planes to a fixed proportion on each bout.

**Figure 5:**
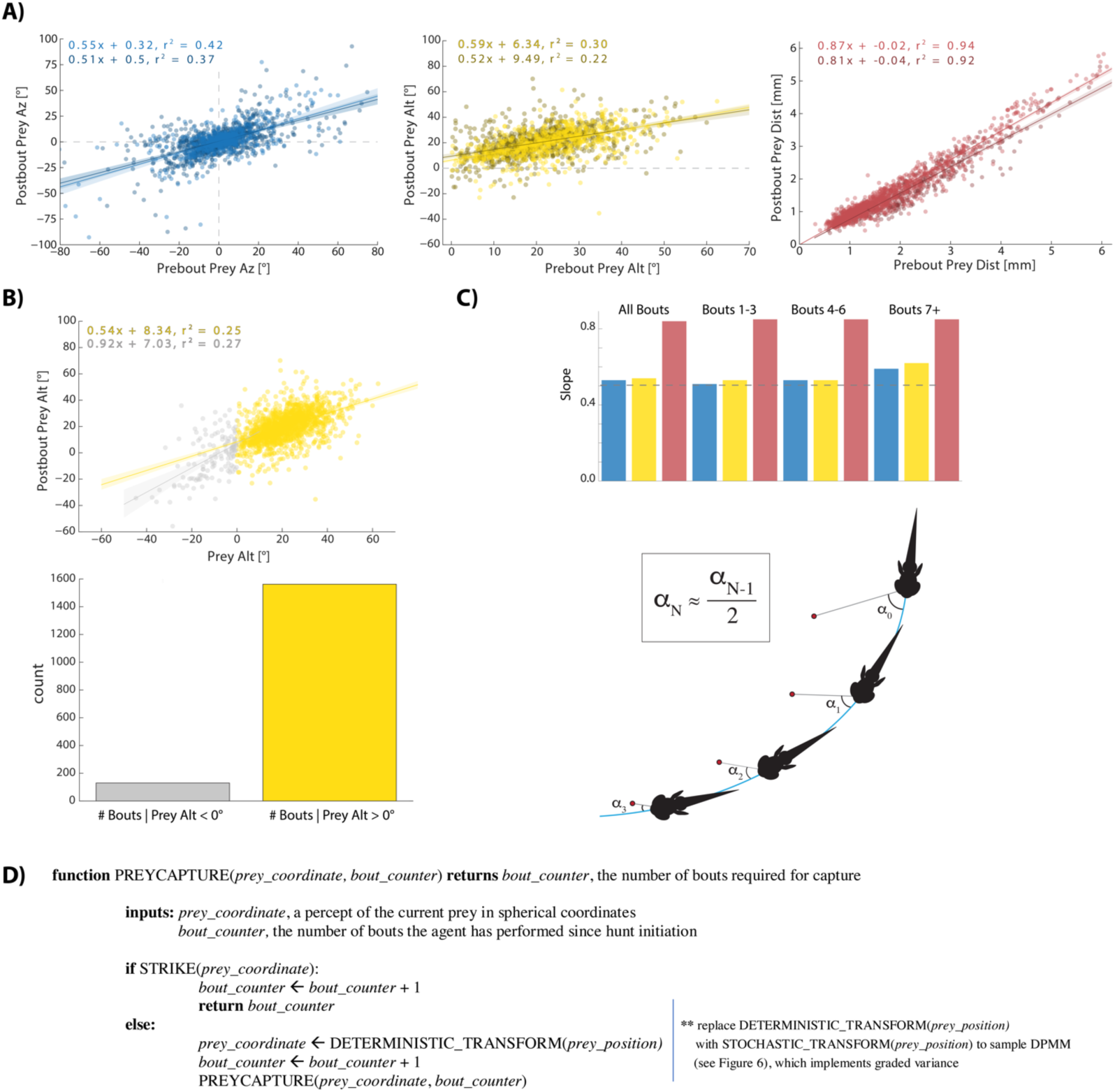
A) Regression plots showing relationships between pre-bout prey coordinates and post-bout prey coordinates. Dark colors, prey is moving towards the fish. Light colors, prey are moving away from the fish. 95% CI on azimuth transforms’ y-intercept includes 0°. B) Regression fits between Pre-Bout and Post-Bout Prey Alt differ depending on whether prey altitude is positive or negative before the bout. Most bouts in the dataset (> 92%) occur when prey are above the fish. C, top) Regression slopes are constant across the hunt sequence; color coded to 5A. C, bottom) Schematic showing recursive halving of the angle of approach during pursuit. D) Pseudocode describing the recursive prey capture algorithm that transforms according to 5A until it arrives at the strike zone.

Interestingly, the .5 slope in Pre-Bout to Post-Bout Prey Alt transformation only holds for when the prey is above the fish (Figure 5B, top). When prey are below the fish, the slope considerably changes to .9, meaning that the fish is only getting 10% closer in altitude per bout. This poor performance for negative Prey Alt is consistent with our observation that hunt initiations are triggered almost exclusively by prey located above the fish. In fact, 92.4% of all bouts in our dataset occur when prey are above the fish (Figure 5B, bottom).

Overall, the fish appears to have a fixed relative 3D angle where it prefers the prey to be at the completion of a pursuit bout. The fact that this location is attained for either direction of velocity is the second piece of evidence that the fish is using velocity projection to estimate future prey position. This “halving” preference is maintained across the duration of hunt sequences, as evidenced by Figure 5C (top panel), which suggests that the fish consistently estimates the future position of the prey before beginning *any* bout. Transforming to a *particular proportion* by abstracting away prey velocity allows the fish to recursively reduce prey coordinates to the termination condition of the algorithm. This is described schematically in Figure 5C (bottom) and Supplementary Figure 5. The combined (both velocity directions) Pre-Bout to Post-Bout Az transform is .*53* * *Pre-Bout Prey Az = Post-Bout Prey Az*. The altitude transformation is .*54* * *Pre-Bout Prey Alt + 8*.*34*° *= Post-Bout Prey Alt*. Implementing these equations iteratively will terminate the algorithm at 18.1° Prey Alt (since .*54* * 18.1° *+ 8*.*34*° *=* 18.1°) and 0° Prey Az, which aligns perfectly with the strike zone described in Figure 2B.

Next, we analyzed how much the radial distance to the prey object is reduced as a consequence of each pursuit bout. We find that on average fish become 16% closer to the prey per bout, or in other words, the Prey Distance is scaled by .84 at every iteration. This reduction, however, is significantly less for prey objects that move radially away from the fish (.87) versus when prey move towards it (.81). This is consistent with the fact that the fish does not modulate Bout Distance based on radial prey velocity when prey are further than 2 mm away. However, once prey are maneuvered to within 2 mm, radial velocity is taken into consideration (Figure 3E). Using radial prey velocity information within 2 mm allows the fish to achieve a fixed Post-Bout Prey Distance, which is reflected by the convergence of the two regression lines in Figure 5A as Prey Distance approaches zero (right panel, dark red and light red). Importantly, 69% of pursuit bouts occur when prey are 0-2 mm from the fish, indicating that fish are most often able to modulate Bout Distance and achieve a preferred Post-Bout Prey Distance after the bout is completed. The combined fit of both velocity directions is .*84* * *Pre-Bout Prey Distance -* .*0125 mm = Post-Bout Prey Distance*.

To summarize, because fish account for prey velocity in all directions (Figure 3), they are on average capable of achieving a fixed proportional reduction of Prey Az, Alt, and Dist during pursuit bouts. Since these proportions are consistent from the beginning to the end of hunting sequences (Figure 5C), we first describe a “deterministic recursive algorithm” for prey capture: The fish recursively transforms current prey coordinates into more favorable prey coordinates by a fixed scale factor in all planes until the strike zone is attained (Figure 5D).

### Graded Stochasticity In Pre-Bout To Post-Bout Prey Coordinates Benefits Hunting Efficiency

We noticed that the variance in pre-bout to post-bout transformation of prey coordinates appeared to decrease with proximity to the strike zone (Figure 5A). In fact, there is a clear graded increase in variance of the post-bout coordinate in all spherical planes as pre-bout coordinates trend away from the strike zone (Figure 6A, left panels). This suggests that the fish is implementing a stochastic recursive algorithm rather than the deterministic recursion described in Figure 5D.

**Figure 6:**
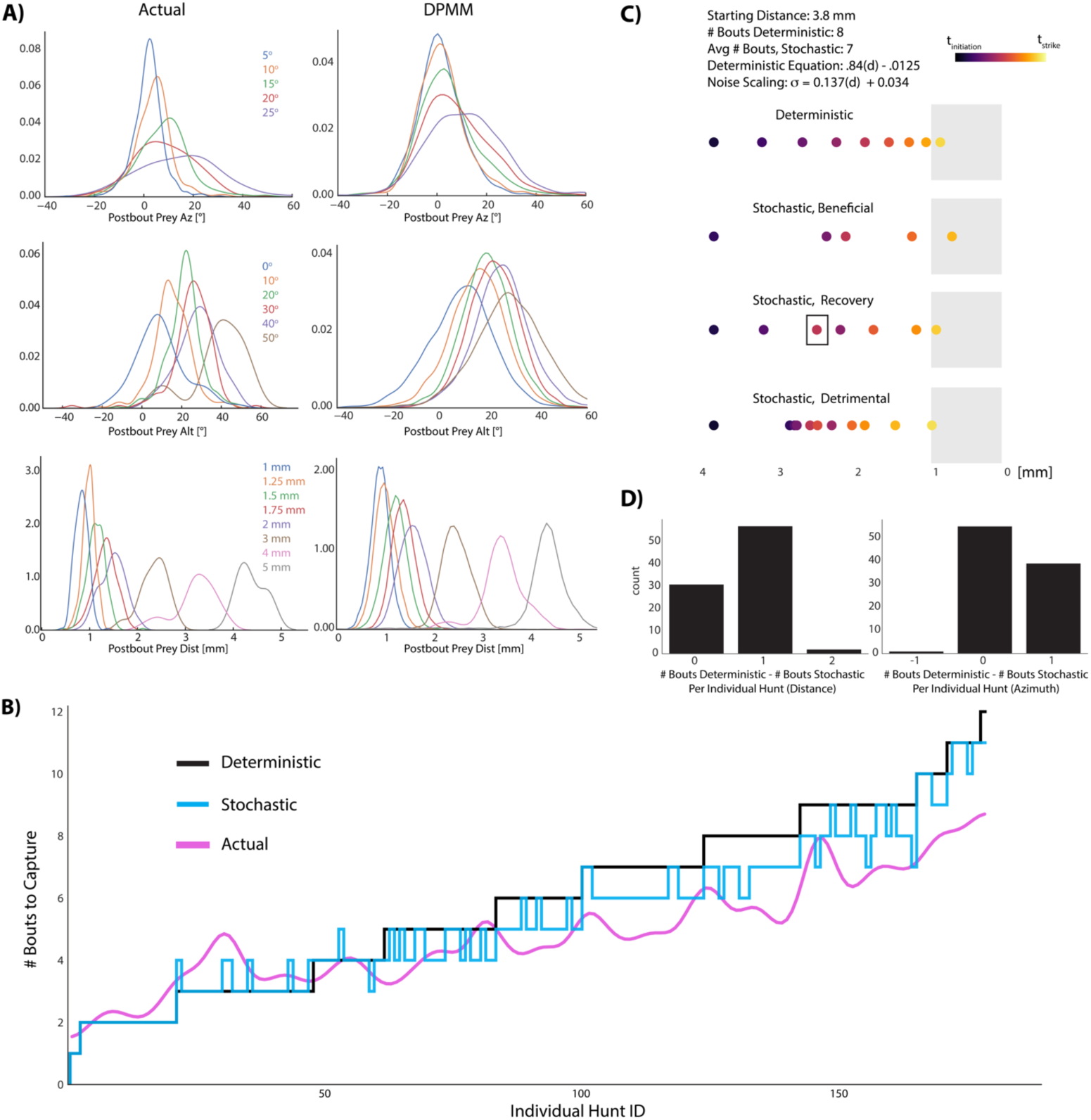
A) KDE plots of post-bout variable distributions for pre-bout input coordinates described in the legend, color-coded to the KDEs (i.e. the blue KDE in Az is the distribution of all Post-Bout Prey Az given that Pre-Bout Prey Az is 5°). Real data is binned in 5° windows for angles and .25 mm bins for distance. DPMM-generated post-bout variables are directly simulated from the model 5000 times, conditioned on the pre-bout value in the legend. B) The median performance of the DPMMs embedded in a recursive loop (stochastic recursion algorithm; run 200 times per initial prey position) typically ties or outperforms the deterministic recursion model transforming with the same average slopes of 10000 pre-bout / post-bout samples generated from the DPMMs. Pink line is the gaussian filtered performance of the real fish (σ = 2) provided the same initial prey condition, showing that the trend of the real fish’s performance closely matches the stochastic recursion. C, D) Graded Variance Algorithm (described in 6C equations) is applied 500 times per initial distance (.1 mm to 10 mm, .1 mm steps) or azimuth (10° to 200° in steps of 2°). Termination condition is a window from .1 mm to 1mm for distance, −10° to 10° for azimuth (see Appendix for full algorithm). Deterministic recursion algorithm is also run on each initial azimuth and distance with .*53* * *az* for azimuth *and* .*84* * *dist -* .*0125 mm* as the fixed transforms. Graded Variance uses these exact transforms as the mean while injecting graded noise: σ_d_ = 0.137 * dist + 0.034 mm ; σ_az_ = .36 * az + 7.62°. (C) shows examples of Graded Variance performance (“stochastic”) vs. deterministic performance. (D) is a barplot comparing the deterministic performance for each input distance and azimuth to the median Graded Variance (“stochastic”) performance.

We next sought to uncover whether a stochastically implemented version of the recursive hunting algorithm was beneficial to the fish. In order to capture the stochasticity of pre-bout to post-bout prey transforms made by zebrafish during pursuit, we chose to use probabilistic generative models (Dirichlet Process Mixture Models: “DPMMs”). Indeed, in order to accurately reflect realistic, stochastic pre-bout to post-bout transformations, our model choice had to be multivariate, heteroskedastic, and include multi-modal probability distributions over pursuit choices. While our linear parametric model captured the average transformation made by the fish in multiple velocity conditions, analytically tractable model families are unable to qualitatively capture the above phenomena. DPMMs can be thought of as “probabilistic programs” that we used to generate conditional simulations of transforms made by the fish given a particular pre-bout prey coordinate (Goodman et al. 2008, Mansinghka et al. 2014, Cusumano-Towner et al. 2019).

Figure 6A (right panels) shows that simulating post-bout prey coordinates from the set of DPMMs for a given pre-bout prey coordinate reveals the same pattern of graded variance observed in the fish. We next fed the deterministic recursive algorithm (Figure 5D) and a stochastic recursive algorithm (DPMM sampling embedded in a recursive loop: see Figure 5D [**]) the 181 initial prey positions that the real fish model captured in our Virtual Prey Capture Environment (Model 1, Blue; Figure 4). Both programs return the number of “bouts” (i.e. pre-bout to post-bout transforms) needed to reach the strike zone. We counted how many bouts the deterministic and stochastic recursive prey capture algorithms took to bring the initial prey coordinates to the strike zone and compared it to the performance of the real fish. Remarkably, the stochastic model outperformed the deterministic model in terms of capture speed, even though both had the same average transformation in all planes (Figure 6B; Wilcoxon signed rank = 7.5 * 10^−17^). This suggested that the fish’s strategy of graded stochastic transforms centered on a preferred post-bout value is actually beneficial versus accurately achieving a fixed, preferred post-bout value. Moreover, although everything except initial prey position has been abstracted away, the performance of the real fish on these initial prey positions matches the stochastic recursive model: the model even seems to recapitulate local fluctuations in the fish’s performance (Figure 6B, pink line).

To assure that graded variance is responsible for the stochastic recursive model’s improvement, we fit a regression model to the standard deviation of Post-Bout Prey Az for given Pre-Bout Prey Az input, and to the standard deviation of Post-Bout Prey Distance for given Pre-Bout Prey Distance (Figure 6C; see standard deviation equation). These regression equations take a pre-bout coordinate as input and return a standard deviation for its post-bout coordinate. We next implemented an algorithm (“Graded Variance Algorithm”) that takes an initial distance or azimuth as input and draws from a gaussian with a mean of the deterministic transform value and a standard deviation determined by the above regression fits (see Appendix, GRADED_VARIANCE algorithm). Again looping these transforms recursively, we input start coordinates and asked how long the recursion would take to reach a fixed strike zone (Figure 6C). The Graded Variance algorithm outperformed the deterministic recursive model for both angle and distance simulations (Wilcoxon Signed Rank: 1.87 * 10^−9^ azimuth, 3.96 * 10^−14^ distance); common scenarios that we observed in these transformations are illustrated in Figure 6C. Although the deterministic algorithm definitively achieves the strike zone in a fixed number of transforms (Figure 6C, top), the Graded Variance algorithm can either directly improve on the deterministic transform (“Beneficial” panel, 6C), start poorly but then recover and outpace the deterministic transforms due to graded noise (“Recovery”, 6C), or perform detrimentally (“Detrimental”, 6C). Nevertheless, the average performance for a given start coordinate is typically equivalent or better by 1 bout (Figure 6D). Given that successful hunts in our dataset had an interquartile range of 4 to 7 total bouts, this improvement by noise constitutes an average 14-25% gain in capture speed. We therefore conclude that the fish’s graded stochasticity produces performance that is curiously beneficial to the fish while hunting prey.

## DISCUSSION

In our study, we uncovered three basic rules that larval zebrafish implement while hunting their fast-moving prey: 1) prey position linearly governs the aspects of five degrees of freedom in which fish can rotate or translate through the water: rotation in yaw and pitch, as well as lateral, vertical and radial displacement; 2) prey velocity modulates all of these aspects of 3D motion and allows the fish to project prey position forward in time; and 3) prey coordinate transformation operates via graded variance based on prey proximity to the strike zone. These three simple rules work in tandem to generate excellent, energy efficient performance in prey capture (Figure 4, 6), which is the most complex behavior that larval zebrafish perform and would appear to require elements of physical knowledge. We focus our discussion on how these rules apply to related studies on the neural mechanisms of prey capture, how examining prey capture at two levels of abstraction was beneficial to our study, and how the fish’s algorithms appear to be built around the inherent constraints of its own body.

### From Algorithms To Neurobiology

With respect to the neural implementations of the algorithms we describe, the transformation of angular prey *position* into informative neural activity is encapsulated to a large extent by the abundant research related to retinotopic maps (e.g. Sperry 1963, Apter 1946, Muto et al. 2013; but see Avitan et al. 2016). The encoding of distance to an object has been well studied in mammalian visual neuroscience and is primarily focused on binocular disparity allowing stereoscopic comparison of each eye’s retinotopic map (DeAngelis et al. 1998), with some studies focusing on monocular motion parallax (e.g. Nadler et al. 2008). Neuronal encoding of object speed, however, requires more sophisticated circuitry and much less is known about its implementation during prey capture. Speed sensitive neurons that discriminate between “fast” and “slow” prey-like stimuli have been uncovered in the zebrafish optic tectum (Bianco and Engert, 2015). The neuroscience of prey capture in the toad has been less directed towards the velocity of the prey and is more focused on spatial attention and configurational detection of worm-like vs. anti-worm-like stimuli (Ewert 1987, Ingle 1975). Ewert discovered that preferences for worm-like artificial stimuli are largely invariant to the stimulus velocity and are much more sensitive to shape and axis of motion; however, he did uncover a set of “small field” tectal neurons that respond to velocity. Other studies in frogs have shown that velocity perception during prey capture is largely used to discriminate between prey types (fast = cricket, slow = termite) in order to choose the correct kinematic capture strategy (Monroy and Nishikawa, 2011). The neurons that differentiate these prey have not been discovered. Unlike these and other reports studying velocity during prey capture (Trivedi and Bollmann 2013, Patterson et al. 2013), we specifically contend that velocity perception is used to point-estimate a future prey position, and that the fish conducts bouts to achieve this estimate, on average, by biasing their prey position-controlled movements. We show that the perception and projection of velocity is key to prey capture success, and that without it, the azimuth and altitude coordinates of prey after bouts are less likely to lie near the strike zone (Figure 4). This type of predictive use of velocity is reminiscent of elegant behavioral studies that have illustrated trajectory prediction in salamanders and dragonflies (Mischiati et al. 2015, Borghuis and Leonardo 2015). Quantitative descriptions of such complex algorithms are an absolute necessity for generating hypotheses about neural implementations (Marr, 1986), and our studies provide the necessary framework to follow this line of inquiry in the context of hunting in larval zebrafish. Virtual prey capture setups for head fixed larvae provide promising inroads for testing these ideas (Bianco and Engert 2015, Avitan et al. 2016, Trivedi and Bollmann 2013).

The stochasticity that gives rise to the graded variance we describe can have several biological sources. It will be important to unravel whether the neural command signals arriving at muscles grow in variance with increased amplitude or if instead the muscles receive fixed input from the brain for a given prey condition, but themselves respond with graded noise. If the source of the variance is largely of a neuronal nature, then it would be interesting to study where exactly in the pathway from sensory areas to motor neurons such noise starts to appear (Stern et al. 2017). Further elucidation of the fish’s probabilistic strategy should eventually integrate our findings into more general theories describing the utility of noisy biological behavior (i.e. stochastic resonance; see Wiesenfeld and Moss 1995, for review) and intentionally probabilistic circuit structures for solving computational problems (Mansinghka, Jonas, and Tenenbaum 2008).

### Strategic Behavior Arises From Simple Behavioral Rules

The stochastic recursive algorithm in Figure 6 describes the progression of pre-bout to post-bout prey coordinates without explicitly accounting for prey velocity or specifically executing fish movements. This transformation pattern, which reveals a preferred future prey position and thus a trajectory prediction ability, emerges from the execution of the position and velocity-based rules described in Figures 2 and 3. Zebrafish prey capture in Figure 6C has, in fact, been reduced to a single input and a recursive series of stochastic divisions with a termination condition, which largely recapitulates the performance of the fish (Figure 6B, DPMM). It is unlikely that we would have found such a straightforward description of proportionality and stochasticity at the lower level of abstraction (i.e. in the fish’s actual sensorimotor transformations), because pre-bout to post-bout prey transformation is a formulation of fish movements along five different axes acting simultaneously. Describing prey capture in this way allowed us to assess the goals of the fish on each bout (i.e. Marr 1986), and may lead to descriptions about how fish are evaluating their own performance during prey capture.

With regards to the benefits of the zebrafish’s strategy, the proportional reduction of angle and distance saves energy at the expense of speed (Figure 4). Ideal bout choice improved speed of prey capture in our modeling data (Figure 4, Model 6); but it did so at the price of spending almost twice as much energy per paramecium captured. This suggests that the increase in feeding rate that the ideal model would afford seems not to be essential for providing an adaptive advantage. Interestingly, the inherent stochasticity in the algorithm does slightly improve the speed of capture (Figure 6) while adding only a modicum of energy expenditure (Figure 4: comparison of Real Fish Model 1 vs. deterministic regression Model 3). This suggests that the fish has evolved a proper balance between energy expenditure and speed of capture. On the whole, the evolution of an efficient algorithm for prey capture in the zebrafish is in agreement with the theme of efficient behaviors arising from simple rules. However, quantifying the fish’s overall energy consumption in a context where they often quit is difficult: energy consumption should therefore be revisited, incorporating work on the decay of the prey capture algorithm in the last 3 pursuit bouts of aborted sequences (e.g. Henriques et al. 2019, Johnson et al 2019).

With respect to hunting schemes, predatory animals have evolved a variety of strategies to optimize pursuit and intercept prey. Tiger beetles, for example, engage in pure pursuit where the angle of attack is kept constant at zero degrees (Haselsteiner et al. 2014). Salamanders, on the other hand, lead the trajectory of their prey (Borghuis and Leonardo 2015). Dragonflies and falcons often utilize a strategy of maintaining a constant line of sight which affords the benefit of motion camouflage (Kane and Zamani 2014, Mizutani et al. 2003; but see Mischiati et al. 2015). Relevantly, dragonflies also implement an implicit predictive model of their prey as well as a model of the effects of their own body movements on prey drift, which foreshadowed the possible use of predictive models across animals with small brains (Mischiati et al. 2015). Larval zebrafish have been assumed to engage in pure pursuit, the simplest and most heuristic of these strategies. However, we find that the strategy used by these animals is more complex and reflects an implicit predictive model of where prey will be at a specified time in the future. Furthermore, the quantal nature of the zebrafish’s swim bouts allowed us to uncover that the angle of attack is recursively and stochastically reduced by an average proportion until the prey enters a terminal strike zone.

### Embodied Physical Knowledge

One branch of artificial intelligence research advocates against a central processing unit where relevant computation occurs in favor of a distributed network of sensorimotor transformations, tuned to the capabilities of the body, that can accomplish the goals of the system (Brooks 1991). Our study reinforces these sentiments and suggests that approaching the study of the brain without considering its embodiment may be precarious.

Specifically, interesting relationships in the data we provide suggest that the fish’s algorithms are built around the capabilities and constraints of its body. First, the amount of “fixed noise” in the fish’s azimuth transformations is 7.64°. This is the standard deviation of Post-Bout Prey Az given a Pre-Bout Prey Az coordinate of 0°, the minimum of graded stochasticity observed in Figure 6A. The standard deviation of the strike zone itself is 7.2° in azimuth. We contend that the similarity between these two numbers suggests that the fish’s strike zone is constructed to deal with noise that the fish cannot overcome in its motor program. If the prey is at, for example, 5° azimuth, performing another bout to get to the very center of the strike zone (∼0° Az) would in many cases worsen the Post-Bout Prey Az coordinate due to fixed noise. Perfection is the enemy of good in this case. In this sense, the command to end pursuit and issue a strike is triggered by a visual releasing stimulus that evolved due to the fish’s own bodily constraints. This is akin to the idea of embodied cognition (Maturana and Varela 1992). Further evidence for embodied knowledge comes from the bias in the fish’s responses to prey altitude. The fish’s algorithm for transforming prey altitude biases the prey to ∼18° above the fish, which aligns almost perfectly with the mean altitude coordinate at which they strike (17.4°, Figure 2, 3, 5). The mechanics of the fish jaw necessitate this: to open its jaw widely for paramecium entrance into the mouth, the fish *must* tilt its head up due to torsional constraints (Mearns et al. 2019). Therefore, the fish’s entire sensation of prey altitude and its method of keeping the prey above it by biasing its bouts downward emerge from the way its jaw co-operates with the rest of its head. Also of note is that graded variance is minimal for altitude transformations at Pre-Bout Alt = 20° (Figure 6A), whereas the azimuth minimum is at 0°; this is also likely a function of its jaw features, which allows the least motor noise when the prey are located in the ideal strike position.

All things considered, the implicit predictive model of 3D prey motion shown in this study is: 1) embodied by the fish’s stochastic recursive algorithm 2) shaped by the constraints and capabilities of the fish and 3) formulated by the interaction of three simple rules. Importantly, these more nuanced features of the fish’s hunting algorithm would not have been revealed without examining prey capture in its more naturalistic 3D setting, which we believe has laid a groundwork for future studies examining the ontogeny, plasticity, and neural implementation of prey capture algorithms and physical knowledge in general (e.g. Avitan et al. 2017).

## METHODS

### Animals

Experiments were conducted according to animal regulations enforced by the NIH and Harvard Institutional Animal Care and Use Committee. All experiments were performed on dpf 7-8 larval zebrafish of the WIK strain. Fish were raised in an automated system where they were delivered paramecia twice a day from dpf4 onward. Importantly, fish in the system experienced a full range of paramecium movement due to the height of the water in their home tank (∼6”). Fish were fasted for 4-6 hrs before experiments.

### Behavioral Setup

After the fasting period, fish were added with ∼100 paramecia (*Paramecia Multimicronucleatum*) in the dark to a 2cm x 2cm x 2cm cube tank made of clear acrylic capped with coverglass. 3.56 megapixel images were simultaneously acquired from the top and side of the tank using two Point Grey Grasshopper 3 NIR cameras triggered by a Pyboard microcontroller at 62.5 Hz. Custom acquisition code was written using C# with the EmguCV library for high-speed video-writing. For the duration of the experiment, fish were illuminated with an infrared LED array, and after 2 minutes in the dark, fish were exposed to a uniform white LED which commenced prey capture. Fish hunted in the white light illuminated condition for 8 minutes before the experiment was terminated. Fish that did not consume more than 1 paramecium over the 8 minute experiment were discarded for analysis (46 / 53 fish passed this criteria).

### Behavioral Analysis

Custom Python software using the OpenCV library was written to extract the body features of the fish (eye convergence angle, tail curvature, yaw, 3D position) and the position of each paramecium in the XY and XZ planes. Pitch was calculated by taking the 2D vertical angle in the side camera and fitting a cone to the fish using the yaw angle from the top camera. (Software is freely available at github.com/larrylegend33.) The tail angle of the fish was fed to a bout detection algorithm that returned frames where swim bouts were initiated and terminated using tail angle variance and bout velocity.

Hunt sequences were identified by spectral clustering (scikit-learn) the continuous eye angle of both eyes over each swim bout for all fish into 5 clusters. One of the 5 clusters showed clear convergence of both eyes at bout initiation (“hunt initiation cluster”), while a second cluster showed clear deconvergence (“hunt termination cluster”, see Supplementary Figure 1A). Custom annotation software cycled through frames marked as hunt initiations and allowed the user to terminate hunt sequences on frames where the fish consumed its prey or clearly quit hunting. Most hunt terminations coincided with the hunt termination cluster.

Upon identifying the frame boundaries of hunt sequences, a 3D prey trajectory reconstruction algorithm was applied that matched prey discovered in the two separate cameras. This is a nontrivial task because the two cameras only overlap in one axis; any two prey items that share similar values in the overlapping plane must be separated using dynamics in time. We therefore matched prey trajectories from the top and side using correlation of velocity profiles and post-hoc 3D position similarity. In this way, each prey item is assigned an ID for a given hunt sequence; the user is required to specify the prey ID that is struck at on strikes, and the “best guess” prey ID that fish pursue during abort sequences. All prey trajectories for each hunt sequence are mapped to a spherical coordinate system based on unit vectors fit from the fish’s XYZ position, pitch, and yaw, with its origin at the fish’s oral cavity. Manual quality control for mistakes in fish characterization or prey reconstruction was applied rarely by eliminating hunt sequences from analysis showing clear mistakes from the computer vision algorithms.

### Regression Fitting and Modeling Environment

All regression models were fit with Generalized Linear Model tools using the Python StatsModels package. When visualized using the Seaborn library in Python, 95% CIs for fits are represented as light shaded regions behind the regression line.

Ideal choice models used in the Virtual Prey Capture Simulation Environment cycled through bouts combined in a “bout pool” from all 46 zebrafish that were performed during sequences that ended in a strike. Each bout during Choice was pre-filtered for bout duration before scoring for prey closeness to the strike zone; ideal bouts could not be shorter than the bout chosen by the fish at that juncture, and could not extend past the time of the next bout chosen by the fish.

Prey trajectories used in the modeling environment were selected from real capture sequences where the fish struck at the prey and the prey was swimming (> 330 microns per second; 89% of all hunted prey records pass velocity criteria, which through inspection distinguishes swimming from floating prey). All virtual hunt reconstructions were initiated with the virtual fish and prey items in the exact same positions and orientations as when the real sequences were initiated. Energy consumption in the virtual environment was calculated under the assumption that the head to center of mass distance for a larval zebrafish is .53 mm (as measured in ImageJ) and the mass of the fish is 1 mg (Avella et al. 2012). Rotational energy of yaw and pitch and kinetic energy of center of mass displacement were added for each bout. Strike zone achievement was defined by the 95% CI on the angular position of a prey item during successful strikes (Figure 2B), likewise conditioned on the radial distance being less than two standard deviations from the mean.

Both regression and ideal models choose initiation bouts and pursuit bouts independently. The first bout of regression models is fit on only initiation bouts, and the first bout of choice models is chosen from the pool of all initiation plus pursuit bouts. This is largely because initiation bout transformations are significantly different from pursuits and likely serve an initial orienting role (Supplementary Figure 4).

### Abstracted Models and Bayesian Nonparametric Methods

Pseudocode describing the transformations of pre-bout to post-bout paramecium locations were written according to the method of Norvig and Russell (2010). We inferred mixture models from empirical data (Pre-Bout Prey Az, Alt, Dist and Post-Bout Prey Az Alt, Dist for all pursuit bouts in the dataset) using a non-parametric Bayesian prior called a Dirichlet Process Mixture Model (DPMM) (Rasmussen 2000, Antoniak 1974, Mansinghka et al. 2016). DPMMs can approximate a broad class of multivariate distributions without requiring a priori specification of the number of components in the mixture model. The mixture models generated via a DPMM prior can be converted to probabilistic programs for inference to generate the kinds of conditional simulations used in Figure 6 (Saad et al. 2019). In this representation, each pre-bout to post-bout prey transformation made by a zebrafish can be thought of as arising from a program that first chooses a prototypical transform (corresponding to a component in the mixture), and then generates a random transform from a distribution over transforms associated with the prototype. We used the BayesDB software library (Mansinghka et al. 2015, Saad et al. 2016) to implement the computations needed to build these models and generate conditional simulations.BayesDB simulations were embedded inside a recursive loop that input an initial prey position and output the number of bouts until striking (see PREYCAPTURE algorithm in Figure 5D with STOCHASTIC_TRANSFORM substitution). When comparing deterministic and stochastic models in Figure 6, the initiation bout for both models was equal and deterministic; only pursuit bouts differed between deterministic and DPMM models.

## ACKNOWLEDGEMENTS

The authors would like to thank Martha Constantine-Paton, Mehmet Fatih Yanik, Misha Ahrens, Rory Kirchner, Rob Johnson, Lilach Avitan, Roy Harpaz, Kirsten Bolton, and Elizabeth Spelke for conversations on the project. Yarden Katz and Hanna Zwaka provided helpful advice on the manuscript. Armin Bahl and Kristian Herrera provided advice and assistance with 3D rendering. This work was funded by a U19 grant from the National Institutes of Health.

## AUTHOR CONTRIBUTIONS

ADB conceived and designed the experimental setup and study in consultation with MH, JJ, FE, and JBT. ADB collected data and wrote analysis code and simulation software in consultation with MH and US; probabilistic models were implemented using the BayesDB framework developed by VKM, US, and FS. FE obtained funding for the project and provided guidance and supervision with JBT and VKM. ADB and FE drafted the manuscript and all authors contributed to the final version.

## REFERENCES

S. Alem, C. J. Perry, X. Zhu, O. J. Loukola, T. Ingraham, E. Søvik, L. Chittka, Associative Mechanisms Allow for Social Learning and Cultural Transmission of String Pulling in an Insect. PLoS Biol. 14 (2016), doi:10.1371/journal.pbio.1002564.

L. Avitan, Z. Pujic, N. J. Hughes, E. K. Scott, G. J. Goodhill, Limitations of neural map topography for decoding spatial information. J. Neurosci. 36, 5385–5396 (2016).

L. Avitan, Z. Pujic, J. Mölter, M. Van De Poll, B. Sun, H. Teng, R. Amor, E. K. Scott, G. J. Goodhill, Spontaneous Activity in the Zebrafish Tectum Reorganizes over Development and Is Influenced by Visual Experience. Curr. Biol. 27, 2407-2419.e4 (2017).

C. E. Antoniak, Mixtures of Dirichlet Processes with Applications to Bayesian Nonparametric Problems. Ann. Stat. 2, 1152–1174 (1974).

J. T. Apter, Eye movements following strychninization of the superior colliculus of cats. J. Neurophysiol. 9, 73–86 (1946).

M. A. Arbib, Levels of modeling of mechanisms of visually guided behavior. Behav. Brain Sci. 10, 407–436 (1987).

M. A. Avella, A. Place, S. J. Du, E. Williams, S. Silvi, Y. Zohar, O. Carnevali, Lactobacillus rhamnosus Accelerates Zebrafish Backbone Calcification and Gonadal Differentiation through Effects on the GnRH and IGF Systems. PLoS One. 7 (2012), doi:10.1371/journal.pone.0045572.

R. Baillargeon, Young infants’ reasoning about the physical and spatial properties of a hidden object. Cogn. Dev. 2, 179–200 (1987).

P. W. Battaglia, J. B. Hamrick, J. B. Tenenbaum, Simulation as an engine of physical scene understanding. Proc. Natl. Acad. Sci. U. S. A. 110, 18327–18332 (2013).

I. H. Bianco, F. Engert, Visuomotor transformations underlying hunting behavior in zebrafish. Curr. Biol. 25, 831–846 (2015).

I. H. Bianco, A. R. Kampff, F. Engert, Prey capture behavior evoked by simple visual stimuli in larval zebrafish. Front. Syst. Neurosci. 5 (2011), doi:10.3389/fnsys.2011.00101.

G. Borghuis, A. Leonardo, The role of motion extrapolation in amphibian prey capture. J. Neurosci. 35, 15430–15441 (2015).

V. Braitenberg, Vehicles: Experiments in synthetic psychology. MIT press; 1986.

H. Brighton, A. L. R. Thomas, G. K. Taylor, D. Lentink, Terminal attack trajectories of peregrine falcons are described by the proportional navigation guidance law of missiles. Proc. Natl. Acad. Sci. U. S. A. 114, 13495–13500 (2017).

R. A. Brooks, Intelligence without representation. Artif. Intell. 47, 139–159 (1991).

S. A. Budick, D. M. O’Malley, Locomotor repertoire of the larval zebrafish: swimming, turning and prey capture. J. Exp. Biol. 203, 2565–79 (2000).

T. S. Collett, Do toads plan routes? A study of the detour behaviour of Bufo viridis. J. Comp. Physiol. □ A. 146, 261–271 (1982).

I. D. Couzin, J. Krause, Self-Organization and Collective Behavior in Vertebrates. Adv. Study Behav. 32, 1–75 (2003).

M. F. Cusumano-Towner, F. A. Saad, A. K. Lew, V. K. Mansinghka, (2019), pp. 221–236.

G. C. DeAngelis, B. G. Cumming, W. T. Newsome, Cortical area MT and the perception of stereoscopic depth. Nature (1998), doi:10.1038/29299.

J. K. Douglass, L. Wilkens, E. Pantazelou, F. Moss, Noise enhancement of information transfer in crayfish mechanoreceptors by stochastic resonance. Nature (1993), doi:10.1038/365337a0.

T. W. Dunn, Y. Mu, S. Narayan, O. Randlett, E. A. Naumann, C. T. Yang, A. F. Schier, J. Freeman, F. Engert, M. B. Ahrens, Brain-wide mapping of neural activity controlling zebrafish exploratory locomotion. Elife. 5 (2016), doi:10.7554/eLife.12741.

E. Gahtan, Visual Prey Capture in Larval Zebrafish Is Controlled by Identified Reticulospinal Neurons Downstream of the Tectum. J. Neurosci. 25, 9294–9303 (2005).

N. D. Goodman, V. K. Mansinghka, D. Roy, K. Bonawitz, J. B. Tenenbaum, in Proceedings of the 24th Conference on Uncertainty in Artificial Intelligence, UAI 2008 (2008).

A. F. Haselsteiner, C. Gilbert, Z. J. Wang, Tiger beetles pursue prey using a proportional control law with a delay of one half-stride. J. R. Soc. Interface (2014), doi:10.1098/rsif.2014.0216.

P. M. Henriques, N. Rahman, S. E. Jackson, I. H. Bianco, Nucleus Isthmi Is Required to Sustain Target Pursuit during Visually Guided Prey-Catching. Curr. Biol. 29, 1771-1786.e5 (2019).

D. Ingle, Focal attention in the frog: behavioral and physiological correlates. Science (80). 188, 1033–1035 (1975).

G. Jensen, Behavioral Stochasticity. Encyclopedia of Animal Cognition and Behavior (2018), pp. 1–5.

E. Johnson, S. Linderman, T. Panier, C. L. Wee, E. Song, K. J. Herrera, A. Miller, F. Engert, Probabilistic Models of Larval Zebrafish Behavior: Structure on Many Scales. bioRxiv, 672246 (2019).

P. Johnson, D. Amso, J. A. Slemmer, Development of object concepts in infancy: Evidence for early learning in an eye-tracking paradigm. Proc. Natl. Acad. Sci. 100, 10568–10573 (2003).

K. P. Körding, D. M. Wolpert, Bayesian integration in sensorimotor learning. Nature. 427, 244–247 (2004).

B. M. Lake, T. D. Ullman, J. B. Tenenbaum, S. J. Gershman, Building machines that learn and think like people. Behav. Brain Sci. 40 (2017), doi:10.1017/S0140525X16001837.

D. Marr. Vision: A computational investigation into the human representation and processing of visual information. Henry Holt and Co., New York, NY 2.4.2 (1982).

HR Maturana, FJ Varela. The tree of knowledge: The biological roots of human understanding. New Science Library/Shambhala Publications; 1987.

V. Mansinghka, P. Shafto, E. Jonas, C. Petschulat, M. Gasner, J. B. Tenenbaum, “CrossCat: A Fully Bayesian Nonparametric Method for Analyzing Heterogeneous, High Dimensional Data” (2015), (available at http://arxiv.org/abs/1512.01272).

V. Mansinghka, E. Jonas, J. Tenenbaum, Stochastic digital circuits for probabilistic inference. Massachussets Institute of Technology Technical Report (2008).

J. C. Marques, S. Lackner, R. Félix, M. B. Orger, Structure of the Zebrafish Locomotor Repertoire Revealed with Unsupervised Behavioral Clustering. Curr. Biol. 28, 181-195.e5 (2018).

D. S. Mearns, J. L. Semmelhack, J. C. Donovan, H. Baier, Deconstructing hunting behavior reveals a tightly coupled stimulus-response loop. bioRxiv, 656959 (2019).

M. Minsky, Society of mind. Simon and Schuster; 1988 Mar 15.

M. Mischiati, H. T. Lin, P. Herold, E. Imler, R. Olberg, A. Leonardo, Internal models direct dragonfly interception steering. Nature. 517, 333–338 (2015).

A. Mizutani, J. S. Chahl, M. V Srinivasan, Insect behaviour: Motion camouflage in dragonflies. Nature (2003), doi:10.1038/423604a.

A. Muto, M. Ohkura, G. Abe, J. Nakai, K. Kawakami, Real-time visualization of neuronal activity during perception. Curr. Biol. 23, 307–311 (2013).

J. W. Nadler, D. E. Angelaki, G. C. DeAngelis, A neural representation of depth from motion parallax in macaque visual cortex. Nature (2008), doi:10.1038/nature06814.

E. A. Naumann, J. E. Fitzgerald, T. W. Dunn, J. Rihel, H. Sompolinsky, F. Engert, From Whole-Brain Data to Functional Circuit Models: The Zebrafish Optomotor Response. Cell. 167, 947-960.e20 (2016).

P. Oteiza, I. Odstrcil, G. Lauder, R. Portugues, F. Engert, A novel mechanism for mechanosensory-based rheotaxis in larval zebrafish. Nature (2017), doi:10.1038/nature23014.

B. W. Patterson, A. O. Abraham, M. A. MacIver, D. L. McLean, Visually guided gradation of prey capture movements in larval zebrafish. J. Exp. Biol. 216, 3071–3083 (2013).

C. E. Rasmussen, The Infinite Gaussian Mixture Model. Adv. Neural Inf. Process. Syst. 12, 554–560 (2000).

D. F. Russett, L. A. Wilkens, F. Moss, Use of behavioural stochastic resonance by paddle fish for feeding. Nature (1999), doi:10.1038/46279.

F. A. Saad, M. F. Cusumano-Towner, U. Schaechtle, M. C. Rinard, V. K. Mansinghka, Bayesian synthesis of probabilistic programs for automatic data modeling. Proc. ACM Program. Lang. 3, 1–32 (2019).

F. Saad, V. Mansinghka, Probabilistic Data Analysis with Probabilistic Programming. Adv. Neural Inf. Process. Syst. 29 (2016) (available at http://arxiv.org/abs/1608.05347).

E. L. C. Shepard, R. P. Wilson, W. G. Rees, E. Grundy, S. A. Lambertucci, S. B. Vosper, Energy landscapes shape animal movement ecology. Am. Nat. (2013), doi:10.1086/671257.

E. Spelke, S. Hespos, Continuity, Competence, and the Object Concept. Lang. Brain, Cogn. Dev. (2018), doi:10.7551/mitpress/4108.003.0027.

R. W. Sperry, Chemoaffinity in the Orderly Growth of Nerve Fiber Patterns. Proc. Natl. Acad. Sci. United States (1963), doi:10.1073/pnas.50.4.703.

S. Stern, C. Kirst, C. I. Bargmann, Neuromodulatory Control of Long-Term Behavioral Patterns and Individuality across Development. Cell (2017), doi:10.1016/j.cell.2017.10.041.

C. A. Trivedi, J. H. Bollmann, Visually driven chaining of elementary swim patterns into a goal-directed motor sequence: a virtual reality study of zebrafish prey capture. Front. Neural Circuits. 7 (2013), doi:10.3389/fncir.2013.00086.

T. D. Ullman, E. Spelke, P. Battaglia, J. B. Tenenbaum, Mind Games: Game Engines as an Architecture for Intuitive Physics. Trends Cogn. Sci. 21 (2017), pp. 649–665.

S. Verma, G. Novati, P. Koumoutsakos, Efficient collective swimming by harnessing vortices through deep reinforcement learning. Proc. Natl. Acad. Sci. U. S. A. (2018), doi:10.1073/pnas.1800923115.

H. R. Wilson, Spikes Decisions and Actions: Dynamical Foundations of Neuroscience (1999).

