## Supplemental Data for "Elements of a stochastic 3D prediction engine in larval zebrafish prey capture"

**SUPPLEMENTAL VIDEO LEGENDS:**

**Supplementary Video 1:** In each video, a hunt is shown from the top and side cameras simultaneously, followed by a virtual reality reconstruction of what the fish is seeing during the hunt. The virtual reality reconstruction, built in Panda3D, is generated from 3D prey coordinates and unit vectors derived from the 3D position, pitch, and yaw of the fish.

**Supplementary Video 2:** Virtual Prey Capture Simulation Environment. Each model from Figure 4 begins at the same position and orientation, and is given the task of hunting the same paramecium trajectory. In this representative hunt sequence, every model except the Random Choice and Multiple Regression (Position Only) models consume the prey (indicated by red STRIKE flash). As is typical in the simulations, the Ideal Choice (Position) model lags the Ideal Choice (Velocity) model by one bout.

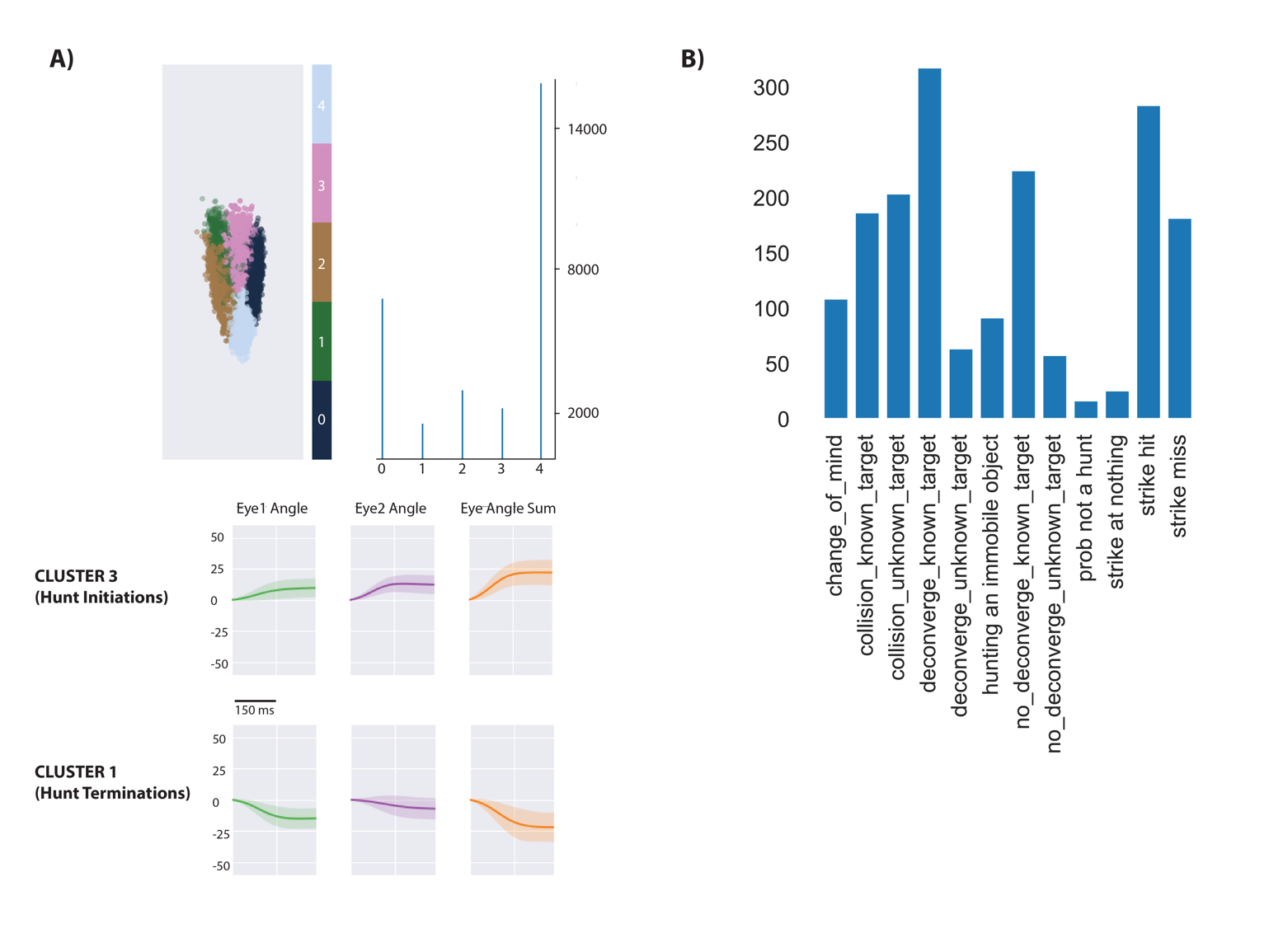

**Supplementary Figure 1:** A) Spectral clustering (scikit-learn) was used to cluster the continuous eye angle over each bout for both eyes. The initiation of hunt sequences was identified using Cluster 3 and deconvergence of the eyes was demarked by Cluster 1. B) Hunt sequence types in the dataset. The user’s only role is to denote the last bout of the hunt sequence, characterize the sequence as a strike hit, strike miss, or abort, and note the chosen prey ID assigned by our automated prey reconstruction algorithm. The program then outputs the descriptors in B per hunt. “Collisions” imply that the fish head has collided with the wall during the hunt (detected using fish COM and edge coordinates), preventing analysis of whether the fish would have struck or aborted. Collisions with unknown target are likely hunting of a paramecium reflection. Deconvergence, known target and unknown target, is the standard abort described previously (Johnson et al. 2019, Henriques et al. 2019), with “unknown targets” being too ambiguous for the user to make a call on pursued prey ID. No deconvergence, known target are hunts where fish had initiated to and pursued a particular prey item, but clearly stopped pursuit on a particular bout not assigned to Cluster 1. “Probably not a hunt” was a rare case where the fish converged, swam through the tank without choosing a prey, and did not deconverge within 8 bouts. “Strike at Nothing” was another rare case where the fish converged and struck without a prey item present. The fish did spend some time striking at immobile objects that were almost invariably residue stuck to the top of the tank. During strike hits, fish choose a prey and consume it, with strike misses typically a deflection of the prey off the fish’s mouth at hunt termination. The main manuscript is built off of strike hits and misses (which combined into a “strikes at known prey” category would be the most common outcome), while Supplementary Figure 3’s abort algorithm is built from known target, deconverge and no-deconverge hunts. Collisions are simply an outcome of having a relatively small tank for parfocal imaging compared to the fish’s real environment; hunts resulting in collisions were not analyzed except for the initial choice.

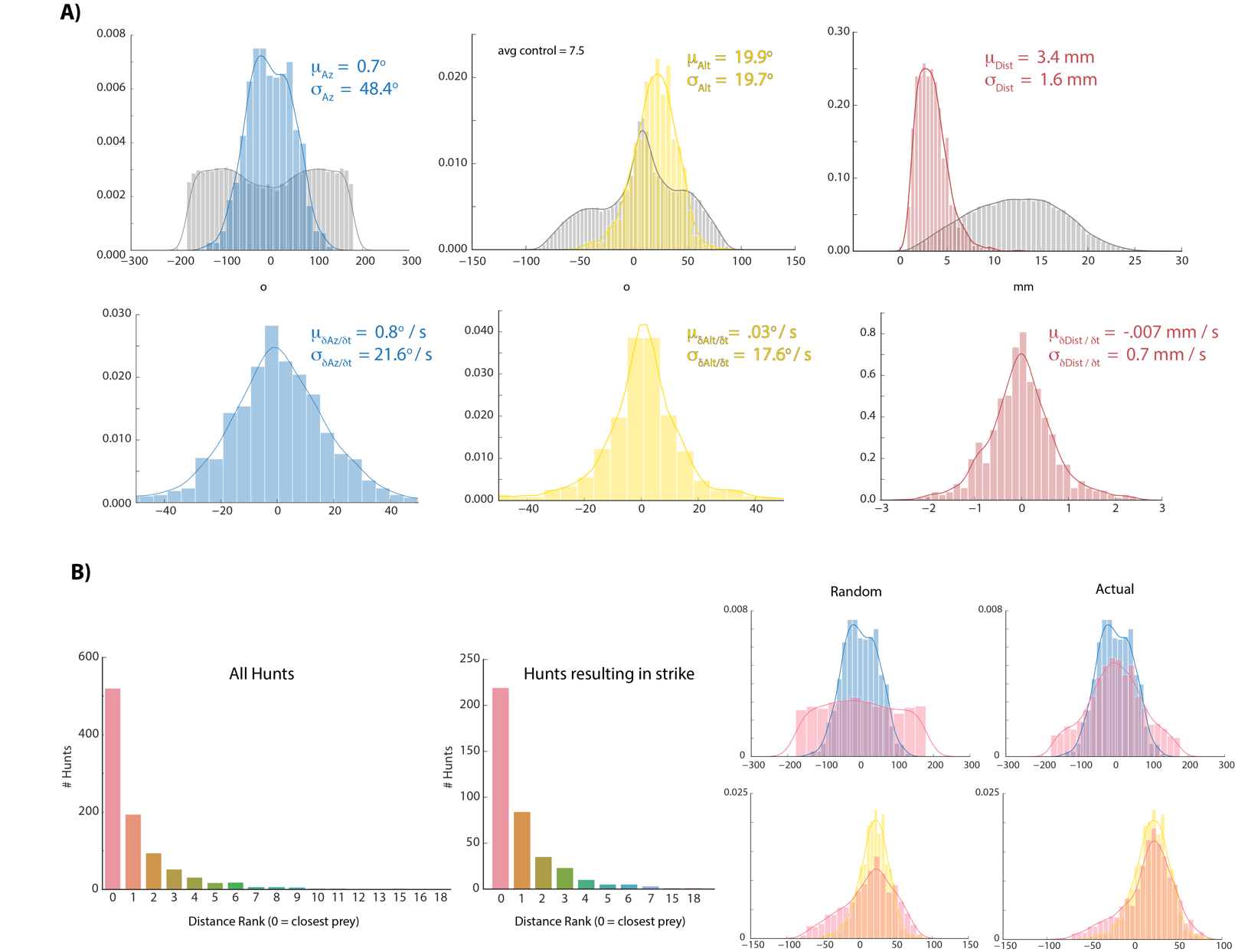

**Supplementary Figure 2:** A) Colors are histograms of coordinates for prey chosen at hunt initiation, gray are all prey records [chosen + ignored. Note: XY records passing a threshold length and velocity that remain unpaired after prey reconstruction are assigned the Z-coordinate of the tank ceiling because live paramecia show anti-gravitaxis (Roberts 2010); we always noted that a subset of prey gather at the ceiling and never noticed coagulation of prey on the ground unless dead; results are very similar to those shown if ceiling assigned prey are not counted]. B) Histograms showing the distribution of spherical velocities for chosen prey do not reveal a bias in magnitude or direction. C) Count plots of distance rank for selected prey (0 = closest) D) We virtually displaced fish coordinates at hunt initiation bouts into randomly recorded paramecium environments and asked whether the closest prey item in that environment shared azimuth and altitude features with prey that fish actually chose. Histograms of the closest prey in *random* prey environments and prey environments in which initiation actually occurred are plotted in red (for random condition, fish orientation and position at hunt initiation is projected into a different time during the experiment; left panel). Blue (az) and yellow (alt) histograms are chosen prey histograms from (A) for comparison. The closest prey item in a random environment does not show the same distribution as selected prey in (A), indicating that the closest prey does not necessarily have to share the altitude and azimuth features of chosen prey. This suggests that somewhat specific prey features are preferred for entry into the hunting state, although transition probabilities governing hunting mode entry are also at play (Johnson et al. 2019, Mearns et al. 2019).

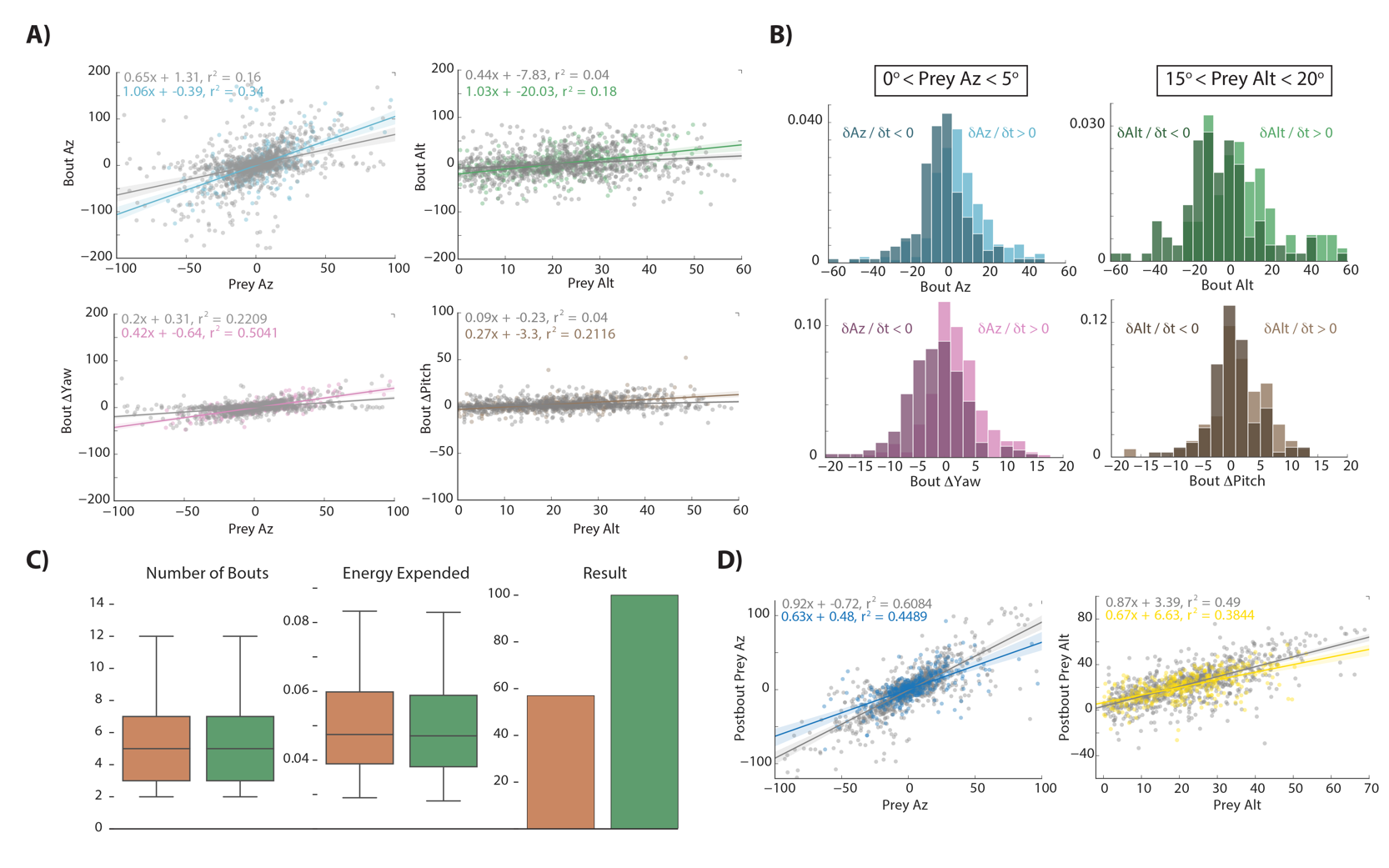

**Supplementary Figure 3:** A) Regression fits between prey position and bout variables during hunt sequences ending in an abort. Gray points and lines represent the last 3 pursuit bouts before the quit bout occurs, colors are all bouts between initiation and the last 3. The algorithm strongly resembles Figure 2 transformations at the beginning of hunt sequences that will eventually end in aborts, but goes awry in the last 3 bouts before quitting. B) Pursuit bouts during abort sequences show modulation by velocity at inflection points similar to Figure 3B. C) Orange model is the same as Figure 4 (Orange Model 2), which issues bouts based on multiple regression to prey position variables only. Green model is same as Figure 4 (Green Model 3), which issues bouts based on multiple regression to prey position and velocity variables. However, both are fit using pursuit bouts during aborted sequences (outside of the last 3 bouts before quitting) instead of strike sequences (i.e. Figure 4). As with models fit on strike sequences, multiple regression using prey position and velocity outperforms position only regression due to proportional velocity modulation. Both models are fed the exact same prey trajectories as models in Figure 4. Fitting on pursuit bouts during aborted sequences thus shows similar performance levels to models fit on strike sequences.

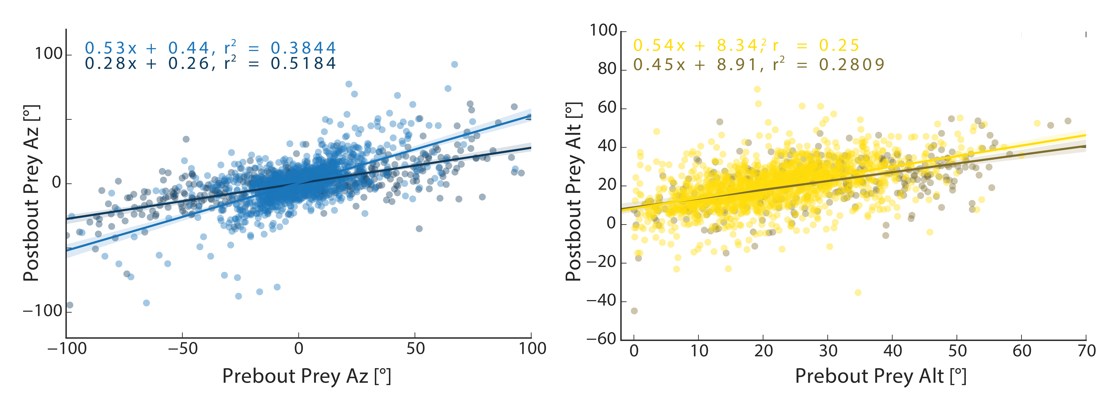

**Supplementary Figure 4:** Transformation by the initiation bout of Pre-Bout to Post-Bout Prey Az and Alt. The initiation bout is a large angle turn that divides azimuth more than a pursuit bout; altitude is also significantly more reduced by the initiation bout. All regression models in simulations therefore use an independent regression fit to initiation bouts to start every simulated hunt.

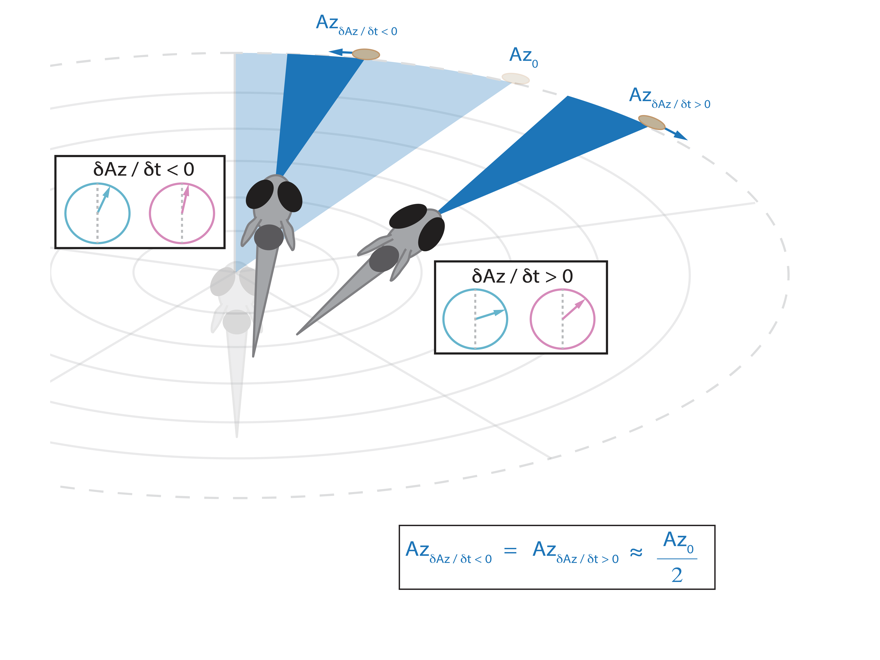

**Supplementary Figure 5:** Schematic showing that the fish will reduce the post-bout angle of attack to the same value regardless of whether prey is moving towards or away from the fish (see Fig 5A). This strategy emerges from the modulation of rotation and displacement by prey velocity (Bout ΔYaw, pink wheel; Bout Az, blue wheel; Figure 3).

**APPENDIX:**

**function** STRIKE(*prey_coordinate*) **returns** *true* if prey in strike zone and *false* otherwise

**inputs:** *prey_coordinate*, a percept of the current prey position

**local variables:** *strikezone*, 95% CI of strike probability based on 2B norm fits, or bounds of a fixed window for single coordinate

**if** *prey_coordinate* in *strikezone* **then return** *true*

**else return** *false*

*­***function** ALT_COEFFS(*prey_alt­­*) **returns** *alt_coefficients,* a list containing a slope and y-intercept for deterministic transform of alt coordinates

**inputs:** *prey_alt*, a percept of the current altitude of the prey item

**local variables:** *alt_slope* | *prey_alt_positive* = .54

*alt_yint* | *prey_alt_positive* = 8.34^o^	from Figure 5B

*alt_slope* | *prey_alt_negative* = .92

*alt_yint* | *prey_alt_negative* = 7.03 ^o^

**if** *prey_alt* > 0, **then return** [*alt_slope* | *prey_alt_positive*, *alt_yint* | prey_alt_positive]

**else return** [*alt_slope* | *prey_alt_negative*, *alt_yint* | *prey_alt_negative*]

**function** DETERMINISTIC_TRANSFORM(*prey_coordinate*) **returns** *new_prey_coordinate,*

**inputs:** *prey_coordinate*, a percept of the current spherical prey coordinate as a list [‘az’, ‘alt’, ‘dist’]

**local variables:** *az_slope* = .53

*alt_slope*

*alt_yint*	from Figure 5A

*dist_slope* = .84

*dist_yint* = -.0125 mm

*new_prey_coordinate,* the new spherical prey position after transform

*alt_slope*, *alt_yint* 🡨 ALT_COEFFS(*prey_position*[‘alt’])

*new_prey_coordinate* 🡨	[*prey_coordinate*[‘az’] * *az_slope*,

*prey_coordinate*[‘alt’] * *alt_slope* + *alt_yint*,

*prey_coordinate*[‘dist’] * *dist_slope* + *dist_yint*]

**return** *new_prey_coordinate*

**function** PREYCAPTURE(*prey_coordinate, bout_counter*) **returns** *bout_counter*, the number of bouts required for capture

**inputs:** *prey_coordinate*, a percept of the current prey in spherical coordinates

*bout_counter,* the number of bouts the agent has performed since hunt initiation

**if** STRIKE(*prey_coordinate*):

*bout_counter* 🡨 *bout_counter* + 1

**return** *bout_counter*

**else: **** replace DETERMINISTIC_TRANSFORM(*prey_position)*

*prey_coordinate* 🡨 DETERMINISTIC_TRANSFORM(*prey_position)* with STOCHASTIC_TRANSFORM(*prey_position*) to sample DPMM

*bout_counter* 🡨 *bout_counter* + 1 (see Figure 6), which implements graded variance

PREYCAPTURE(*prey_coordinate*, *bout_counter*)

**function** GRADED_VARIANCE(*prey_coordinate, bout_counter, dist_or_angle*) **returns** *bout_counter*, the number of bouts required for capture

**inputs:** *prey_coordinate*, a percept of the current prey position in spherical coordinates

*bout_counter,* the number of bouts the agent has performed since hunt initiation

*dist_or_angle*, string representing whether input is a distance or an azimuth angle

**local_variables:**

μ, a value representing the average transform

σ, the standard deviation of the average transform that decreases with proximity to the strike zone

**if** STRIKE(*prey_coordinate*):

*bout_counter* 🡨 *bout_counter* + 1

**return** *bout_counter*

**else:**

**if** *dist_or_angle* == ‘angle’:

μ 🡨 53 * *prey_coordinate*

σ 🡨 .36 * *prey_coordinate* + 7.62°

**if** *dist_or_angle* == ‘distance’:

μ 🡨 84 * *prey_coordinate* - .0125 mm

σ 🡨 0.137 * *prey_coordinate* + 0.034 mm

*prey_coordinate* 🡨 GAUSSIAN_DRAW(μ, σ)

*bout_counter* 🡨 *bout_counter* + 1

GRADED_VARIANCE(*prey_coordinate, bout_counter, dist_or_ang***)**
